# FastCAT accelerates absolute quantification of proteins by using multiple short non-purified chimeric standards

**DOI:** 10.1101/2021.12.28.474379

**Authors:** Ignacy Rzagalinski, Aliona Bogdanova, Bharath Kumar Raghuraman, Eric R. Geertsma, Lena Hersemann, Tjalf Ziemssen, Andrej Shevchenko

**Affiliations:** Max Planck Institute of Molecular Cell Biology and Genetics, 01307 Dresden, Germany; Center of Clinical Neuroscience, Department of Neurology, University Hospital Carl Gustav Carus, Technical University of Dresden, 01307 Dresden, Germany

**Author notes:** Corresponding author: Dr. Andrej Shevchenko.

**Keywords:** absolute quantification of proteins, MS Western, QconCAT, targeted quantitative proteomics, cerebrospinal fluid, neurodegeneration, neuroinflammation

## Abstract

Absolute (molar) quantification of clinically relevant proteins determines their reference values in liquid and solid biopsies. FastCAT (for <u>Fast</u>-track Qcon<u>CAT</u>) method employs multiple short (<50 kDa) stable-isotope labeled chimeric proteins (CPs) composed of concatenated quantotypic (Q-) peptides representing the quantified proteins. Each CP also comprises scrambled sequences of reference (R-) peptides that relate its abundance to a single protein standard (BSA). FastCAT not only alleviates the need in purifying CP or using SDS-PAGE, but also improves the accuracy, precision and dynamic range of the absolute quantification by grouping Q-peptides according to the expected abundance of target proteins. We benchmarked FastCAT against the reference method of MS Western and tested it in the direct molar quantification of neurological markers in the human cerebrospinal fluid at the low ng/mL level.

## INTRODUCTION

The role of absolute (molar) quantification of proteins is multifaceted. It determines stoichiometric ratios within molecular assemblies and metabolic pathways^1^ and relates them to the abundance of non-proteinous compounds *e*.*g*. enzyme co-factors, lipids or metabolites. It also provides reference values and ranges of their physiological variation for diagnostically important proteins in liquid and solid biopsies.^2^ Last but not least, it estimates the proteins expression level in cells and tissues serving as a quantitative denominator common to all *omics* sciences. In contrast to popular immunodetection methods (*e*.*g*., ELISA or Western blotting),^3^ mass spectrometry quantifies proteins by comparing the abundance of endogenous quantotypic (Q-)peptides with corresponding synthetic peptide standards having the exactly known concentration. The protein concentration is then inferred from the concentrations of Q-peptides. However, protein and peptide properties are unique and may vary significantly.^4^ Furthermore, to support clinical diagnostics, it is often necessary to quantify a selection of disease-related proteins whose molar abundance differs by several orders of magnitude. It is therefore not surprising that, in contrast to the relative quantification, absolute quantification methods lack generality and unification.

Absolute quantification (reviewed in^5^) relies upon different types of internal standards, including (but not limited to) synthetic peptides (*e*.*g*., AQUA)^6^; full-length or partial protein sequences (*e*.*g*. PSAQ^7^ and QPrEST^8^, respectively) and also chimeric proteins composed of concatenated quantotypic peptides from many different proteins (QconCAT)^9^ (reviewed in^10– 13^). Powered by the recent advances in gene synthesis, QconCAT offers several appealing qualities such as ease of multiplexing that enables targeted mid-to large-scale quantification of individual proteins, protein complexes, metabolic pathways^14^ or selections of clinically relevant of proteins.^15,16^ QconCAT chimeras are expressed in *E*.*coli*, enriched and purified by affinity chromatography and their stock concentration is determined by amino acid analysis or some protein assays^17^. Alternatively, an additional (secondary) peptide concatenated standard (PCS) could help to quantify multiple primary PCSs.^18^ Although cell-free expression systems^19,20^ improve the flexibility of QconCAT implementation, it does not alleviate the need to enrich, purify and quantify the CPs. The sequence of [Glu^1^]-Fibrinopeptide B could be included as a reference, however CP quantification based on a single synthetic peptide standard should be used with caution.^21^

By using GeLC-MS/MS, MS Western workflow alleviated the need in making and standardizing a purified stock of CP. Also, CP standards were designed such that they included not only quantotypic (Q-)peptides, but also several reference (R-)peptides.^22^ The bands of CP and of the reference protein (BSA) were co-digested with gel slabs containing target proteins and the recovered peptides analyzed by LC-MS/MS. By using R-peptides, the abundance of CP was referenced *in-situ* to the exactly known amount of BSA, which is available as a NIST certified standard. Next, the abundance of target proteins was calculated from the abundance of CP assuming that its complete tryptic cleavage produced corresponding Q-peptides in the equimolar amount. MS Western quantification relied upon the concordant values (CV <10 %) obtained from multiple (usually 2 to 4) Q-peptides per each protein of interest and took advantage of high expression of CP in *E*.*coli*.^23,24^

By using SDS for proteins solubilization, GeLC-MS/MS could detect more membrane proteins. SDS-PAGE also alleviated the need in purifying CPs, yet it limited the analyses throughput. While assembling hundreds of Q-peptides into a large (up to 290 kDa) chimera is appealing, it is also inflexible because other proteins and/or peptides could not be added at will. Furthermore, the yielded Q-peptides are strictly equimolar, which hampers the quantification of proteins having drastically (more than 100-fold) different in their abundance. This, however, is often required for quantifying protein biomarkers.^25^

Here, we report on FastCAT (for **Fast**-track Qcon**CAT**) method that preserves the accuracy and consistency of MS Western quantification, yet it is faster, more flexible and easier to use particularly in translational proteomics applications. In contrast to MS Western, FastCAT workflow relies on the parallel use of many relatively short (less than 50 kDa) non-purified CPs that comprise Q-peptides for many target proteins, but also scrambled R-peptides to reference each CP concentration to the same BSA standard.

## EXPERIMENTAL SECTION

### Chemicals and reagents

LC-MS grade solvents (water, acetonitrile and isopropanol), formic acid (FA) and trifluoroacetic acid (TFA) were purchased from Thermo Fisher Scientific (Waltham, MA) or Merck (Darmstadt, Germany). Trypsin and trypsin/Lys-C proteases (MS grade) were from Promega (Madison, WI); RapiGest detergent from Waters (Eschborn, Germany); other common chemicals from Sigma-Aldrich (Munich, Germany). Polyacrylamide gradient gels (4-20 %) were from Serva Electrophoresis GmbH (Heidelberg, Germany). Protein standards: glycogen phosphorylase (GP), alcohol dehydrogenase (ADH), enolase (ENO) and ubiquitin (UBI) were purchased as a lyophilized powder from Sigma-Aldrich. Reference protein standard (BSA, Pierce grade, in ampoules) was from Thermo Fisher Scientific (Waltham, MA). Isotopically labeled amino acids (^13^C_6_,^15^N_4_-L-arginine (R) and ^13^C_6_-L-lysine (K)) were purchased from Silantes GmbH (Munich, Germany).

### Design and expression of chimeric protein standards

In total, six CP standards of different molecular weights (MW) were designed and expressed. General scheme of CP design is presented in Figure S1, while other details including MWs, isotopic enrichment labeling efficiency and full-length sequences are in Table S4. DNA sequences encoding CPs were codon-optimized for *E*.*coli* by GenScript on-line tool and synthesized by GenScript (Piscathaway, NJ). CPs were produced using pET backbone (Novagenn) and *E*.*coli* BL21 (DE3) (argA lysA) strain auxotrophic for arginine and lysine supplemented with ^13^C_6_,^15^N_4_-L-arginine and ^13^C_6_-L-lysine as described.^22^ *E*.*coli* strain^26^ was a kind gift from Professor Roland Hay (University of Dundee, UK).

### Sample preparation for proteomics analyses

Polyacrylamide gels were stained with Coomassie CBB R250 and gel slabs corresponding to the targeted range of MW were excised. In-gel digestion with trypsin^22^ with enzyme-to-substrate ratio of 1:50. *E*.*coli* proteins with 4 spiked standard proteins was in-pellet digested with trypsin (1:20) after protein precipitation with isopropyl alcohol.^27^ Aliquots of cerebrospinal fluid (CSF) of 20 µL volume were in-solution digested with trypsin/Lys-C protease mix (1:20) in the presence of RapiGest (Waters) detergent.^28^ CSF samples were obtained from patients diagnosed with multiple sclerosis and stored as freshly frozen aliquots. All patients gave their prior written consent. The study was approved by the institutional review board of the University Hospital Dresden (EK348092014).

### LC-MS/MS analyses

LC-MS/MS was performed on a Q Exactive HF (Thermo Scientific, Germany) hybrid tandem mass spectrometer coupled with Eksigent 400 nanoLC system (Sciex, Germany) using the Nanospray Flex source (Thermo Fisher Scientific, Germany). Protein digests were loaded onto a trap column for 5 min at 7 µL/min and separated on the Acclaim PepMap 100 column (C18, 3 µm, 75 um x 150 mm) using 120 min gradients (5-45% B) at 300 nL/min in data-dependent acquisition (DDA) or parallel reaction monitoring (PRM) modes (as specified). DDA and PRM methods consisted of MS1 scan from *m/z* 350 to 1700 with automatic gain control (AGC) target value of 3×10^6^; maximum injection time (IT) of 60 ms and targeted mass resolution R_*m/z*=200_ of 60,000. Top-12 DDA method employed the precursor isolation window of 1.6 Th; AGC of 1×10^5^; maximum IT of 50 ms; R_*m/z*=200_ of 15,000; normalized collision energy (NCE) of 25% and dynamic exclusion of 30 s. The scheduled PRM method acquired MS/MS using 10-min retention time (RT) windows with the inclusion list of 99 precursors; precursor isolation window of 1.6 Th; AGC of 1×10^6^; a maximum IT of 80 ms; R_*m/z*=200_ of 30,000 and NCE of 25%.

### Data processing and analysis

Raw LC-MS/MS data from DDA experiments were processed by Progenesis LC-MS v.4.1 (Nonlinear Dynamics, UK) software for RT alignment, peak picking and extracting the peptide features. To match peptides to target proteins, MS/MS spectra were searched by Mascot v.2.2.04 software (Matrix Science, London, UK) against a customized database containing sequences of all target proteins and relevant (either *E*.*coli* or human) background proteome. Precursor mass tolerance of 5 ppm; fragment mass tolerance of 0.03 Da; fixed modification: carbamidomethyl (C); variable modifications: acetyl (protein N-terminus) and oxidation (M); labels: ^13^C_6_(K) and ^13^C_6_^15^N_4_ (R), cleavage specificity: trypsin with up to 2 missed cleavages allowed). All PRM data sets were analyzed with Skyline 21.1.0.278 software.^29^ Peak integration was inspected manually. Mass transitions in labeled and unlabeled peptides and matching retention time and peak boundaries were confirmed the peptide identities. A minimum of five transitions were required for the correct identification of the targeted peptides. In addition, the comparison of the measured fragment spectrum to the *in-silico* Prosit-derived^30^ library spectrum by the normalized spectral contrast angle was used that resulted in the library dot product (dotp) correlation values of 0.85 or higher.

## RESULTS AND DISCUSSION

### Using crude CP standards for protein quantification

Trypsin cleavage of a purified CP produces Q- and R-peptides in a strictly equimolar concentration.^22,31,32^ However, in a crude extract of expression host cells (*E*.*coli*) the concentration balance between individual Q-peptides could be affected by the non-proportional contribution of products of intra-cellular proteolysis of CP and/or incomplete translation of the CP gene. Chimeric proteins are highly expressed in *E*.*coli* and are spiked into the analyzed sample in a minute (low picomole to femtomole) amount^22^. While CPs purification should reduce the concomitant load of *E*.*coli* proteins, it is unclear if their contribution to overall compositional complexity is substantial.

Therefore, we set out to test if full-length CP standards could be spiked directly as crude *E*.*coli* extracts with no prior purification. To this end, we selected three CP standards spanning a wide MW range (CP01 ∼265 kDa, CP02 ∼79kDa and CP03 ∼42 kDa) (Supplementary Table S4). We loaded protein extracts onto 1D SDS-PAGE, excised the CP bands, and also sliced the entire gel slab in several MW ranges below the band of CP and analyzed them separately by GeLC-MS/MS (Figure 1). We observed that the relative abundance of Q-peptides in each slice depended on molecular weight of the full-length CP and its truncated forms, but also on Q-peptides location within the CP sequence. In 265 kDa CP peptides located closer to its N-terminus were overrepresented and constituted 40 to 80 % of total peptide abundances (Figure 1A). We also observed the same trend for the middle-size CP with 20 to 40 % fraction of truncated forms and slight prevalence of N-terminally located peptides (Figure 1B). However, in the shortest (42 kDa) CP the contribution of truncated forms was minor (ca 10-15 %) and independent of peptides location (Figure 1C).

**Figure 1.**
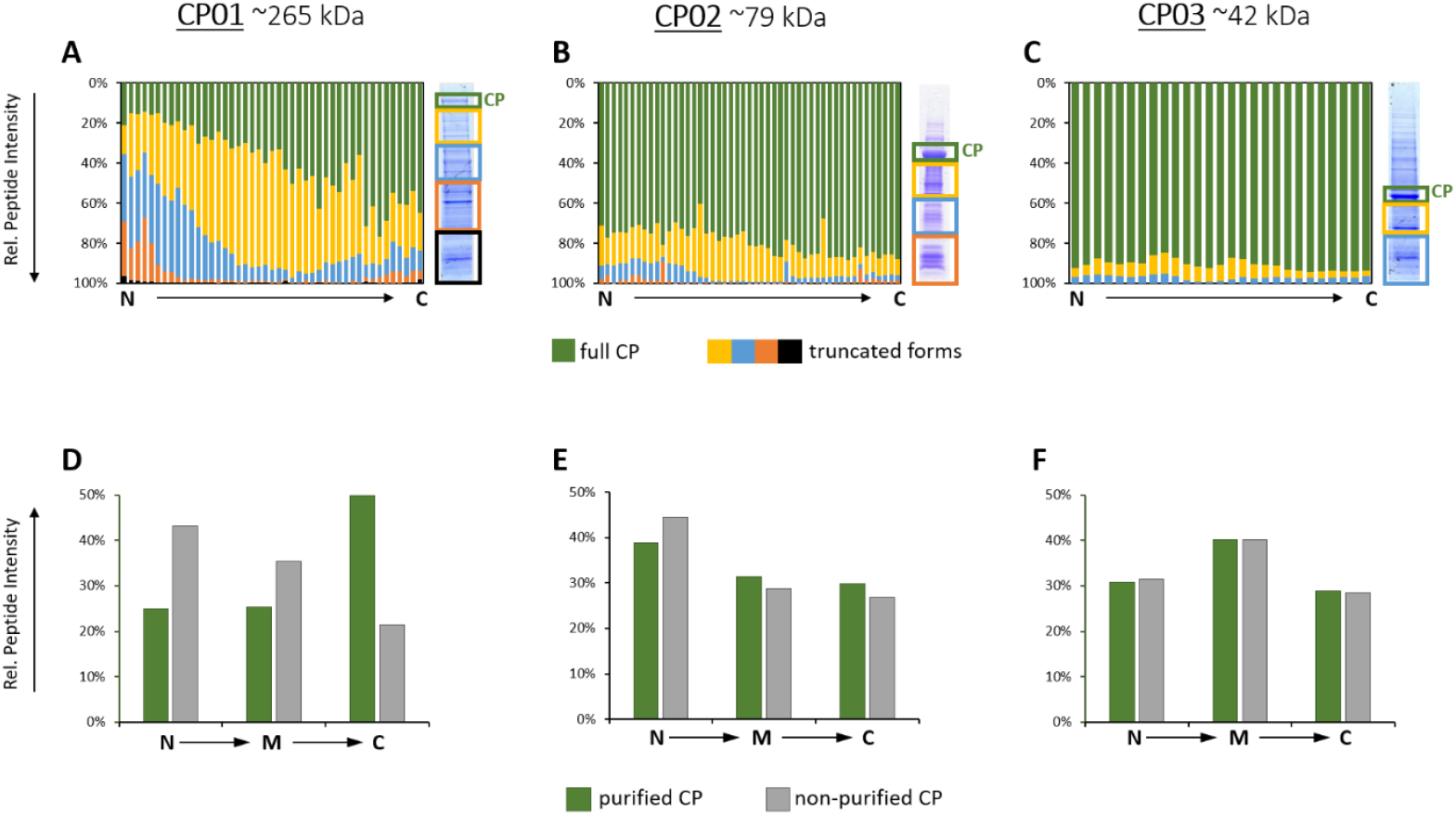
Truncation patterns for the chimeric proteins: CP01 ∼265 kDa (panels: **A, D**), CP02 ∼79 kDa (panels: **B, E**), CP03 ∼42 kDa (panels: **C, F**). The upper panels (**A, B, C**) present relative abundance of peptides in SDS-PAGE slabs (*y*-axis) *versus* peptides position in the CP sequence (*x*-axis); color-coding is exemplified at the right-hand side panel. Lower panels (D, E, F) present relative abundance of peptides from the purified CP (only CP band) versus the non-purified CP (sum of all bands) for selected Q-peptides located at the N- and C-termini as well as in the middle of the CP sequence (“M”).

Because of lower expression and higher fraction of proteoforms having incomplete sequences truncated elsewhere at the C-terminal side, larger CP required extensive purification. In contrast, shorter (less than *ca*. 50 kDa) CPs are highly expressed and cell lysate contained much lower fraction of truncated forms with no location prevalence. Consistently, the relative abundances of Q-peptides at different locations within the CP sequence (N-terminal *vs*. middle (“M”) *vs*. C-terminal) in the purified CP and non-purified CP (CP band together with truncation products) were very close (Figure 1E). Therefore, we concluded that spiking the total *E*.*coli* extract without isolating the full-length CP should not bias the quantification.

Since short CPs are highly expressed in *E*.*coli* such that they are the most abundant proteins in a whole lysate (Supplementary Figure S2), the lyzates are usually diluted more than 100-fold down to *ca*. 1 µM (or even lower) concentration of CP. Therefore, we wondered if adding the extract of *E*.*coli* containing an appropriate amount of non-purified CPs still elevates the protein background. For reliable comparison, we mixed a volume of metabolically labeled extract containing typical working amounts of CP (*ca*. 100 fmol) with an equivalent volume of unlabeled extract containing *ca*. 500 ng of total protein. We then compared relative abundances of labeled and unlabeled forms of nine major *E*.*coli* proteins (Supplementary Table S1) and observed that adding an extract with unpurified CP increased protein background by as little as 2 % (Supplementary Figure S3).

We therefore concluded that short ∼50 kDa CPs could be spiked into quantified samples as total (crude) *E*.*coli* lysates with no prior purification. With that size, they would be typically encoding for 25 to 30 Q-peptides and 3 to 5 R-peptides within a typical CP construct.^22^

### FastCAT workflow: the concept and its validation

We reasoned that, by employing short non-purified CP, a targeted absolute quantification workflow (termed FastCAT for **Fast**-track Qcon**CAT**) could significantly accelerate the analysis. Besides Q-peptides used to quantify target proteins, a typical FastCAT construct contains multiple (usually 3 to 5) R-peptides used to determine the *in-situ* CP concentration by referencing it to the known amount of spiked-in BSA. Hence, FastCAT workflow requires neither CP purification (externally as a stock or by using the band from gel electrophoresis) nor separate determination of its concentration, while the multi-peptide quantification procedure is the same as in MS Western.

We cross-validated FastCAT by comparing it against MS Western.^22^ To this end, we prepared an approximately equimolar (*ca*. 1 µM) mixture of the 4 standard proteins (GP, UBI, ADH and ENO) and spiked it (*ca*. 20 pmol of each protein) into *E*.*coli* background (*ca*. 50 µg of total protein). We determined their exact quantities by MS Western using 42 kDa (CP03) as an internal standard.^22^ In parallel, we quantified them by the FastCAT protocol using the same 42 kDa construct, but without 1D SDS-PAGE. Importantly, we checked the digestion completeness for both methods by comparing relative abundances of labeled Q-peptides in CP03 and corresponding unlabeled peptides in the endogenous proteins. The difference in protein quantities determined by FastCAT and MS Western were below 15 % for all proteins (Table 1).

**Table 1.**
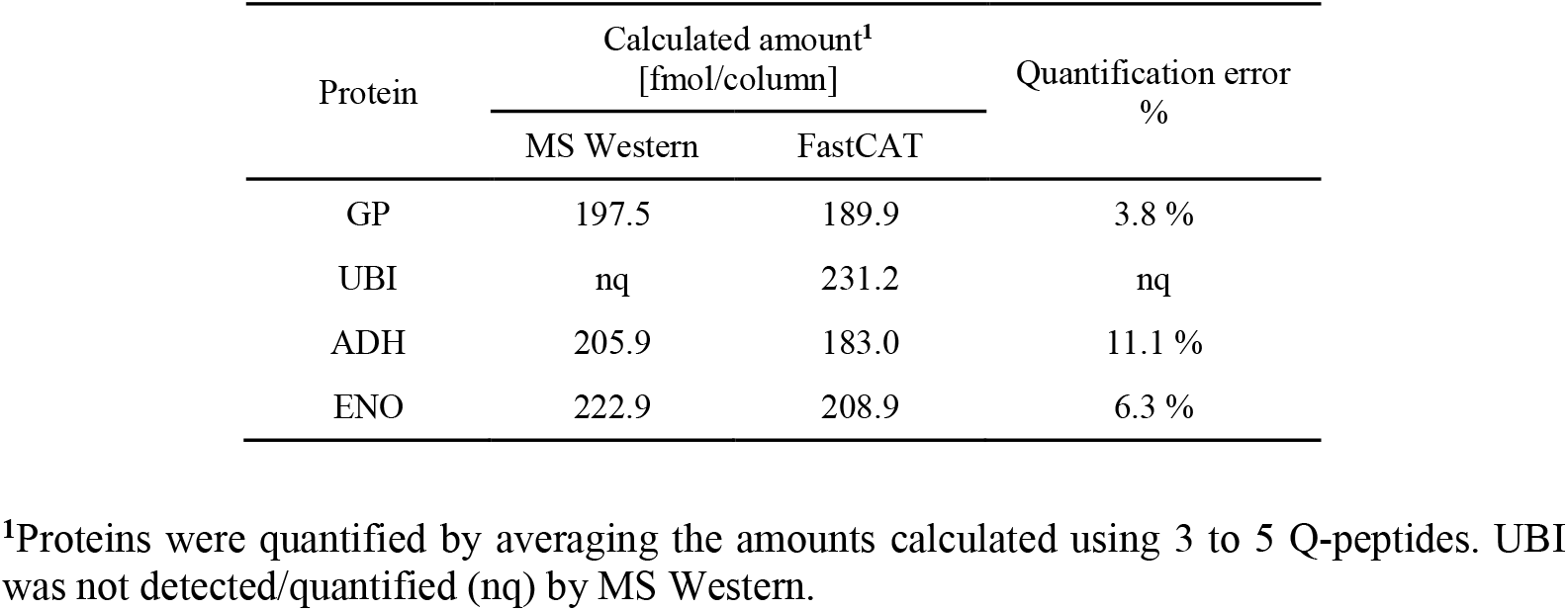
Comparison of protein quantitation by FastCAT and MS Western.

Since in FastCAT workflow requires no purification of CPs, we additionally checked if the positioning of Q- and R-peptides within the CP backbone affected the quantification. To this end, we designed another CP standard, CP04, having the same Q-peptides as in CP03, but in which the same five R-peptides (R1-R5) were distributed over the entire CP sequence, in contrast to a single block of R-peptides at the C-terminus of CP03 (see Figure 2).

**Figure 2.**
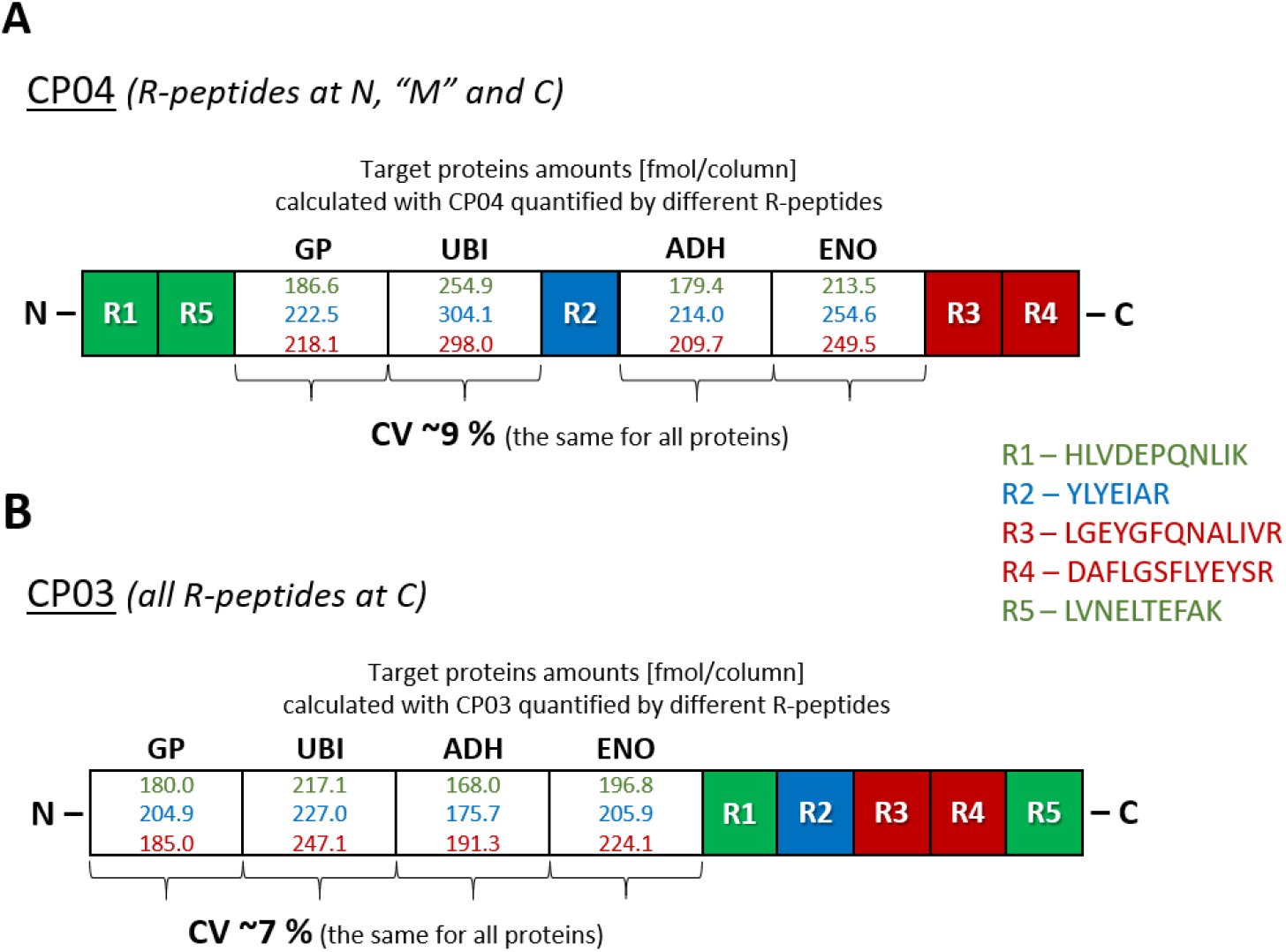
Impact of the position of R-peptides within CP sequence on proteins quantification. Comparison of the target proteins quantification by using CP04 and N-*vs*. “M” *vs*. C-terminus R-peptides as well as by using CP03 and the same groups of peptides but all positioned at C-terminus.

This experiment resulted in two major findings. Firstly, using CP03 and CP04 independently led to concordant quantification. With all R-peptides used for calculating CPs abundances, the quantities of target proteins differed by less than 15%. Secondly, the quantification was practically unaffected by positioning of R-peptides. Indeed, using differentially positioned R-peptides (N (R1/R5) *vs*. “M” (R2) *vs*. C (R3/R4)) in CP04 led to the quantification of all target protein with CV *ca*. 9 % (Figure 2A). This relatively minor variability could not be solely attributed to R-peptides placement. We note that the value of CV *ca*. 7 % was observed when target proteins were quantified using CP03 with R-peptides (R1/R5 *vs*. R2 *vs*. R3/R4) placed at the C-terminus, as was in CP04 (Figure 2B).

We therefore concluded that in the FastCAT workflow the design of CP and relative positioning of R-peptides have no major impact on the proteins quantification.

### Multiplexing of FastCAT

Relatively short CP standards having MW of 40 to 50 kDa will typically comprise 20 to 30 Q-peptides. Considering that the robust protein quantification requires 3 to 5 peptides per each protein, one CP should enable the quantification of 5 to 8 individual proteins of, preferably, similar abundance. Hence, there is a clear need in multiplexing the FastCAT quantification capacity.

We therefore proposed to group proteotypic peptides into CPs according to the expected abundance of the target proteins and then to use multiple CPs in parallel to eventually cover the desired number of proteins (Figure 3). However, the abundance of different CPs should be referenced to the same spiked-in BSA standard. We achieved it by including reference peptides with the scrambled sequences (R_s_-peptides)^33^ that, nevertheless, elicit very similar response in MS1 spectra compared to corresponding native R_n_-peptides. However, we reasoned that, for better sensitivity, the targeted analysis would require PRM also for quantifying CPs in the same LC-MS/MS run. Therefore, for each R_n_/R_s_ peptides pair, we selected a representative combination of fragment ions that adequately reflected the peptides abundance^34,35^ and, hence, enable parallel quantification of multiple CPs.

**Figure 3.**
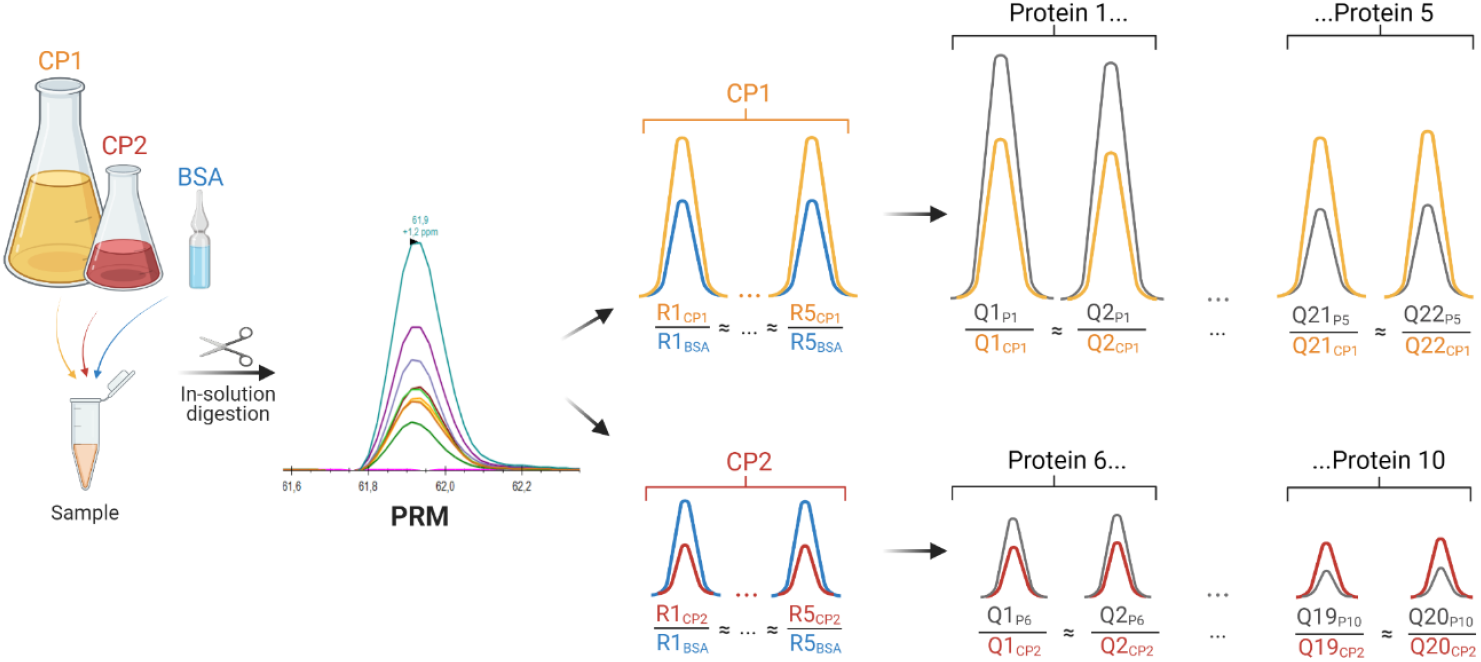
FastCAT workflow for the absolute quantification of proteins. Target proteins (Protein 1, …, Protein 10) having a broad range of molar concentrations are quantified using two unpurified chimeric standards CP1 and CP2. CP1 contains quantotypic peptides from P1 to P5 proteins (Q1_P1_, Q2_P1_, … to …Q21_P5_, Q22_P5_); while CP2 – from the P6 to P10. The amount of CP1 and CP2 spiked into the sample is adjusted to match the expected range of proteins concentration, while the concentration of reference protein (BSA) is known. The sample and spiked proteins are digested and analyzed in PRM mode. First, CP1 and CP2 concentrations are determined by comparing their “heavy” native or scrambled reference peptides (R1_CP1_ … R5_CP1_; R1_CP2_ … R5_CP2_) and matching native (R1_BSA_ … R5_BSA_) peptides from BSA. Then, the concentration of each protein is calculated from peak areas of endogenous Q-peptides (Q1_P1_, Q2_P1_, … to Q21_P5_, Q22_P5_) and matching “heavy” Q-peptides (Q1_CP1_, Q2_CP1_, …, Q21_CP1_, Q22_CP1_). Proteins whose Q-peptides are comprised in CP2 are quantified similarly. Altogether, each protein is independently quantified by at least 2 Q-peptides and referenced to the known amount of BSA. Since peptide concentrations are equimolar protein quantities should corroborate.

To this end, we designed CP05 and CP06 proteins (see Supplementary Figure S4 for amino acid sequences and Figure S5 for the distribution of their truncated forms) containing 42 Q-peptides from 10 selected human proteins (these CPs were further used in the case study described in the next section). They also comprised 10 R-peptides as 5 pairs of native (R_n_) and scrambled (R_s_) sequences. However, R-peptides were placed into CP05 and CP06 such that they contained 3 native plus 2 scrambled, and 2 native plus 3 scrambled R-peptides, respectively (Figure 4). To emulate the impact of protein background, we spiked these CPs in *ca*. 1 µM concentration into a 20 µL aliquot of a human cerebrospinal fluid (CSF) and analyzed their tryptic digests by the method of PRM.

**Figure 4.**
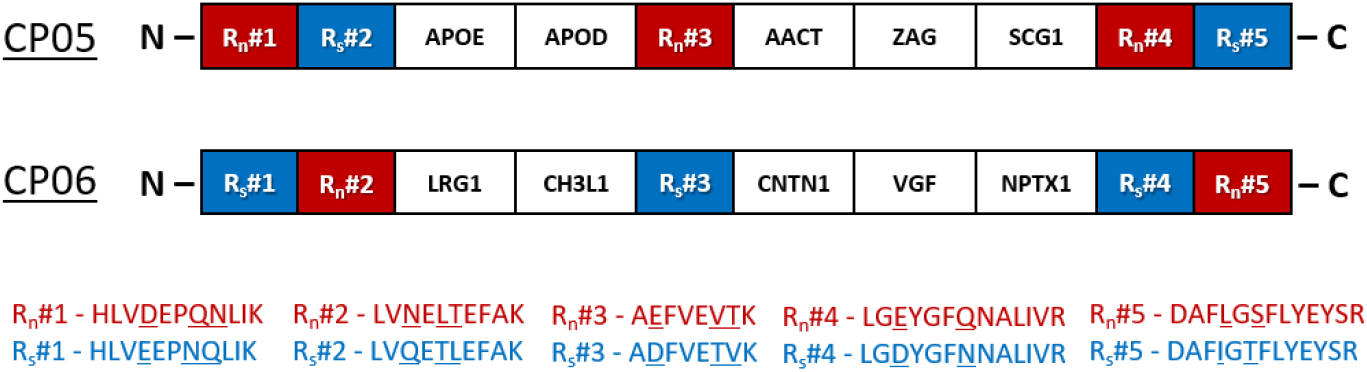
Scheme of CP05 and CP06 constructs comprising the native (R_n_#1 – R_n_#5) and scrambled (R_s_#1 – R_s_#5) forms of 5 BSA peptides (shown as one block/peptide) as well as the multiple Q-peptides from 10 target proteins (shown as one block/protein).

For each R_n_/R_s_ peptide pair we compared the distribution of abundances of the most intense fragments within y- and b-ions (see Supplementary Figure S6 for corresponding MS2 spectra). Out of 5 pairs, three pairs produced very similar profiles, while two other pairs mismatched (Figure 5).

**Figure 5.**
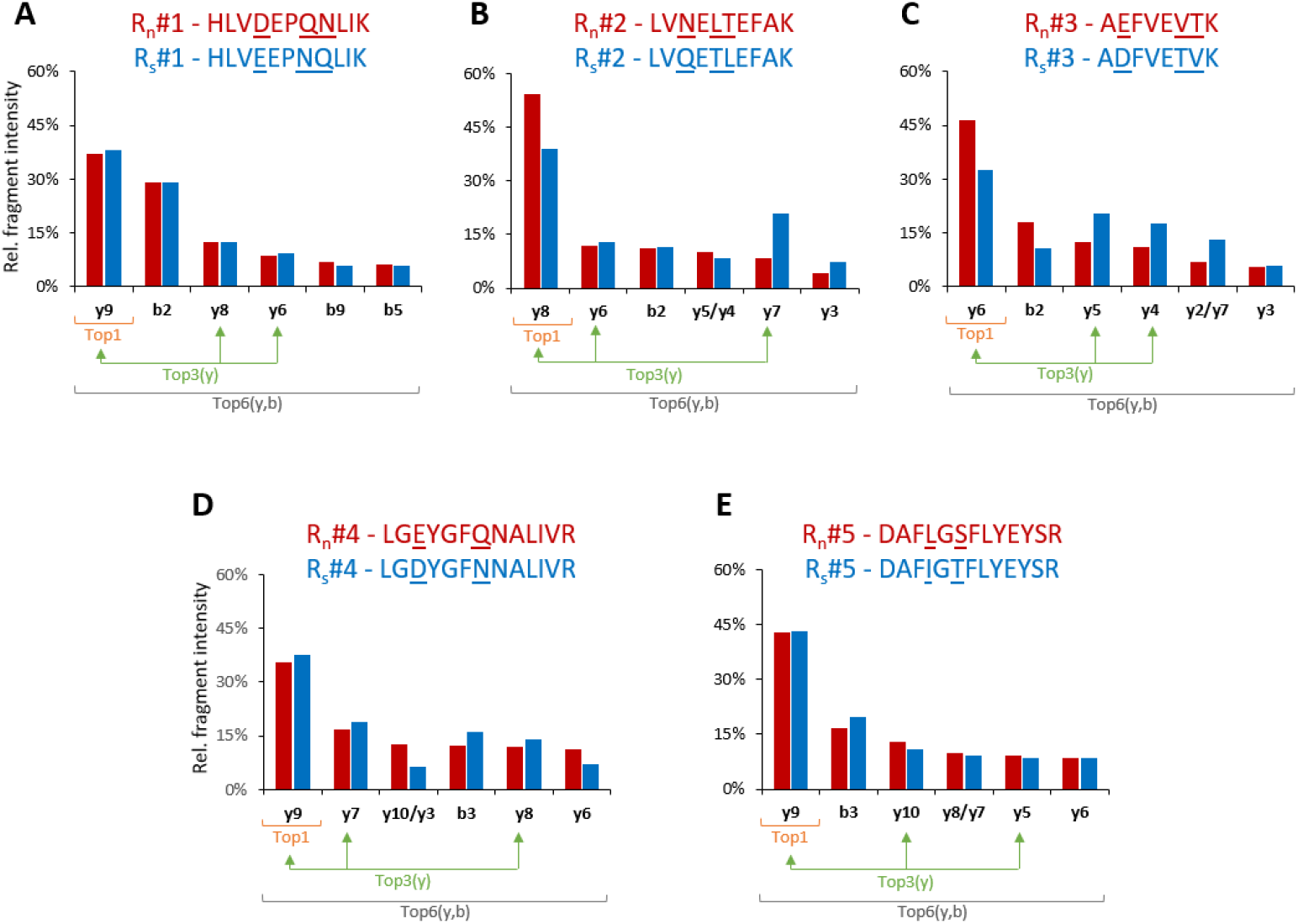
Relative abundance of the most intense six fragment ions (within y- and b-ions) from the native (BSA/light – red bars) and scrambled (CP/heavy – blue bars) R-peptides. In four of five pairs one fragment ion within the top 6 did not match between the native and scrambled forms and in this situation the non-matching fragments were presented together for better visibility of the fragmentation differences.

Next, we assessed the concordance of the *in-situ* PRM-based quantification of CPs using each R_s_-peptide and R_n_-peptides. To do this, we considered three PRM quantification scenarios based on the selection of different combinations of fragment ions: Top1, Top3(y) and Top6(y,b) (as exemplified in Figure 5); and compared them to the values computed from the abundance of MS1-peaks (see Supplementary Table S2). As expected, the Top1 approach not only resulted in the highest error (>35 %) for the two mismatching R_n_/R_s_ (#2 and #3) pairs, but also revealed the highest overall (for all R_n_/R_s_ pairs) discordance with MS1 (Figure 6). In contrast, summing up the abundances of more fragments in Top3(y) or Top6(y,b) scenarios compensated minor differences in the fragmentation patterns of R_n_/R_s_-peptides. We note that for selected reaction monitoring (SRM) experiments performed on low mass resolution instruments that cannot follow many transitions in parallel and where using b-ions should be avoided,^36^ the Top3(y) approach could be the most practical compromise.

**Figure 6.**
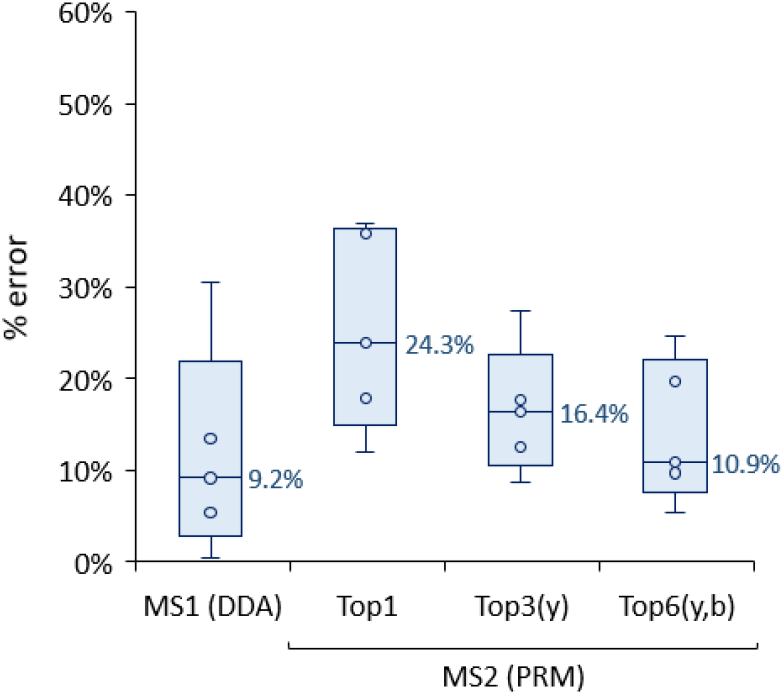
Relative error in the quantification of CPs by using R_s_-peptides and either MS1 or MS2 (summed) intensities of different fragment ions. Top1 (the top fragment ion); Top3(y) (sum of the top three y-ions, matching between R_n_ and R_s_); Top6(y,b) (sum of the top six fragments within y- and b-ions). Each box contains information from 5 R_s_-peptides (detailed information is provided in Table S2 in SI), while the median values are given next to the boxes. Each box plot displays the median (line), the 25th and 75th percentiles (box), and the 5th and 95th percentiles (whiskers).

However, for PRM-based quantification summing up the abundance of 6 most abundant y- and b-fragment ions will lead to the most consistent estimates with lower than 20 % difference to MS1-based determinations (Figure 6). Eventually, within the two CPs the relative abundances of all used (both native and scrambled) R-peptides measured via Top6(y,b) fragments differed by less than 5% (Supplementary Figure S7).

We therefore concluded that FastCAT quantification can be multiplexed by simultaneously using several CPs comprising multiple scrambled peptides of BSA. Quantities of individual CPs could be referenced to the same BSA standard also by the PRM analysis that relies on the summation of six most intense y- and b-ions for each R_s_ (in the CP) and R_n_ (in BSA standard) peptide pair.

### Case study: absolute quantification of neurological protein markers in the human CSF

Multiple sclerosis is an immune-mediated demyelinating and neurodegenerative disease of the central nervous system (CNS), which is accompanied by blood-brain barrier disruption, infiltration of immune cells into the CNS, nerve fibers demyelination and axonal loss.^37^ It alters the CSF proteome and monitoring the levels of protein markers by common clinical chemistry methods (*e*.*g*. ELISA) aids in its molecular diagnostics.^38^ However, these methods suffer from low concordance, limited scope and substantial costs. Here we employed FastCAT to determine the molar concentration of a selection of protein markers having a broad range of physicochemical properties, molar abundance and magnitude of response towards the disease.

We obtained 11 samples of CSF from five (four female and one male) 30 to 61 years of age patients that were diagnosed with relapsing-remitting multiple sclerosis. CSF was drawn from each patient at two time points: the first puncture was performed during the initial diagnostics, while the second (and, for one patient, also the third) puncture was performed *ca*. 2 years later prior to a planned treatment switch to validate that no significant inflammation and neurodestruction occurred. Based on clinical indications, ten protein markers were selected out of *ca*. 700 proteins detected in a pooled CSF sample by the preliminary experiment. Those included two major lipoproteins (APOE and APOD); inflammation-related glycoproteins (AACT, ZAG and LRG1); markers of axonal (CNTN1) and synaptic (NPTX1 and VGF) related disorders; a member of the granins family (SCG1) and, finally, a neuroinflammatory marker (CH3L1) typically increased in patients with multiple sclerosis.^39,40^ We then selected 42 Q-peptides and assembled them in CP05 and CP06 (both mentioned above) according to the arbitrary abundance of target proteins.

The method precision was evaluated by processing and analyzing the pooled CSF sample in triplicate (Figure S7). For both CP and target proteins peptide and protein levels the median coefficient of variation was below 6 %. Importantly, peptides originating from the same protein led to their highly concordant quantification as exemplified by median CV of *ca*. 12 %.

The median values and the ranges of variation of target proteins are reported in Table 2, which also include Q-peptides and the estimates of concordance for the independent quantification by multiple peptides. Concentration ranges determined by FastCAT corroborated previously reported SRM and PRM determinations. For instance, similar concentration ranges were obtained for APOE,^41,42^ AACT,^43^ SCG1^44^ and CH3L1.^45^ At the same time, the concordance with ELISA measurements was limited for both FastCAT and published SRM/PRM values. While ranges determined by ELISA for AACT,^46^ LRG1,^47,48^ CH3L1^45,49,50^ and CNTN1^46^ were close to FastCAT, levels for both apolipoproteins were discordant (yet, again, concordant with SRM/PRM).^51–54^ This, however, is consistent with known discrepancy between ELISA and mass spectrometry measurements.^41^ Molar concentration of ZAG, VGF and NPTX1 was not reported previously.

**Table 2.**
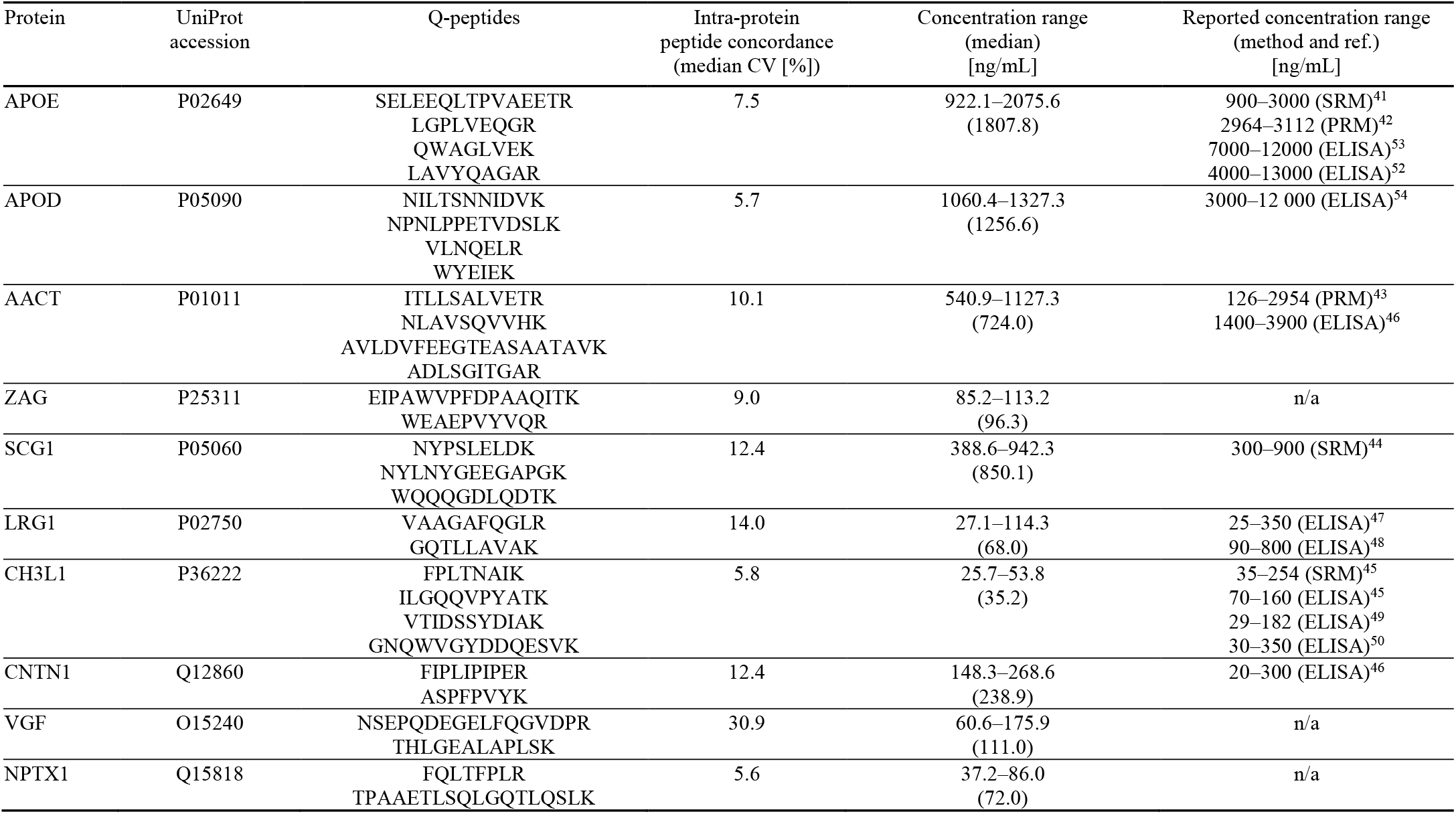
Proteins concentration (ranges and median) determined in eleven CSF samples from patients with multiple sclerosis together with UniProt accession numbers; Q-peptides used for the quantification; intra-protein concordance (median CV from all samples) as well as the previously reported concentrations determined by mass spectrometry and/or ELISA. Proteins: APOE (apolipoprotein E), APOD (apolipoprotein D), AACT (Alpha-1-antichymotrypsin), ZAG (Zinc-alpha-2-glycoprotein), SCG1 (Secretogranin-1/Chromogranin B), LRG1 (Leucine-rich alpha-2-glycoprotein 1), CH3L1 (Chitinase-3-like protein), CNTN1 (Contactin-1), VGF (Neurosecretory protein VGF), NPTX1 (Neuronal pentraxin-1).

Concentrations determined in individual patients (Supplementary Table S3) were stable over the two-years period of treatment and, except one patient, clustered together at the PCA plot (Supplementary Figure S9). PCA plot singled out one female patient presumably because of her higher age, although proteins concentrations in both of her biopsies were concordant.

Several trends (*e*.*g*. increase in APOD and ZAG; decrease in NPTX1) corroborated previous reports.^55–57^ At the same time, there was no consistent change in the levels of prospective multiple sclerosis markers SCG1 and LRG1.^48,58^

Notably, the amount of spiked CP05 standard was ca 10-fold higher than of CP06 (309 fmol/column *vs*. 33 fmol/column, respectively). For better consistency, BSA was spiked at some intermediate amount (100 fmol/column). In this way, PRM covered 100-fold range of concentrations from *ca*. 20 ng/mL for CH3L1 to *ca*. 2000 ng/mL for APOD without breaking inter-peptide quantification consistency. Considering the observed signal-to-noise ratios, PRM was not close to the limit of detection and, in principle, should allow us to reach 10-fold higher sensitivity if the appropriate amount of yet another CP standard is spiked into CSF samples. Taken together, we demonstrated that FastCAT supported direct molar quantification of 10 neurological protein markers is CSF at the low ng/mL levels, better that 100-fold dynamic range and good (CV <20 %) inter-peptide quantification consistency based on two to four peptides per each protein.

## CONCLUSION AND PERSPECTIVES

Absolute quantification of proteins offers heavily missing data on the molar concentrations (or molar abundances) of proteins in cells, tissues and biofluids. FastCAT workflow takes advantage of the flexibility and ease of use of peptide-based quantification. It relies on a large number of peptide standards having the exactly known and equimolar concentration that are produced *in-situ* and quantified in the same LC-MS/MS run together with peptides from target proteins.

In this work we used two CPs to quantify 10 neurological markers in the human CSF. However, FastCAT offers ample capabilities for multiplexing. First, re-engineering reference peptides combined with alternative metabolic labeling (*e*.*g*. by using different combinations of commercially available isotopologues of arginine and lysine) could produce dozens of R-peptides to accommodate in many more unique CPs. A rough calculation suggests that we might only need to design five more R-peptide variants to employ 10 CPs in parallel. If each 50 kDa CP comprises 3 Q-peptides per target protein, this would already cover 100 proteins in a single run. Modern analytical solutions such as SureQuant or PRM-Live offer intelligent real-time PRM scheduling and enable the quantification of hundreds of peptide precursors without compromising the sensitivity and throughput. Furthermore, PRM quantification could be organized in a “modular” fashion by combining CPs of the desired composition and similar concentration range. CP standards are produced by expressing synthetic genes in *E*.*coli* and, because of consistently high expression levels (Supplementary Figure S2), could be used directly from a host cell lysate without their prior purification.

In the future, it might be practical to set up a publicly available repository of plasmids encoding CPs. This will improve the analyses consistency and, eventually, bring absolute quantification availability and performance closer to clinical chemistry requirements. While the throughput of LC-MS/MS quantification could hardly compete with ELISA, its accuracy, independence of antibodies, quantification transparency and analytical flexibility, including compatibility with major protocols for biochemical enrichment and robotic sample preparation, might be appealing for translational applications. We also argue that the interested laboratories should work together towards benchmarking and validating the absolute quantification methods by ring trials that are now common in neighboring omics fields, *e*.*g*. lipidomics.^59^

## Supporting information

FastCAT_Supporting Information

## SUPPORTING INFORMATION

The following supporting information is available free of charge at ACS website http://pubs.acs.org.

Figure S1. Scheme of chimeric protein (CP).

Figure S2. Gel images of short CPs.

Figure S3. *E. coli* background in non-purified CP.

Figure S4. Sequences of CP05/CP06.

Figure S5. Truncation patterns for CP05/CP06.

Figure S6. Tandem mass spectra for Rn/Rs-peptides.

Figure S7. Relative Rn/Rs-peptides abundances for CP05/CP06 vs. BSA.

Figure S8. Precision of FastCAT method.

Figure S9. PCA plot for CSF proteins from patients.

Table S1. E. coli background proteins.

Table S2. Error in quantification of CPs by Rs-peptides.

Table S3. Protein concentrations in CSF.

Table S4. Sequences for all CPs.

## Author Contributions

I.R. and A.S. conceptualized and designed FastCAT. I.R. performed the experiments and interpreted the data. A.B. and E.R.G. produced chimeric proteins. T.Z. provided the CSF samples and clinical support. B.K.R. provided expert technical support. I.R. and A.S. wrote the manuscript.

## Notes

The authors declare no competing financial interest.

Raw DDA data have been deposited at the ProteomeXchange Consortium via the PRIDE partner repository with the dataset identifier PXD030728. PRM data processed by Skyline are available at Panorama Public at https://panoramaweb.org/FastCAT.url.

## ACKNOWLEDGMENTS

Work in Shevchenko group is supported by Max Planck Society. The authors are grateful for other group members for expert technical support and useful discussions. Figures were assembled using the BioRender tool from BioRender.com.

## For TOC only

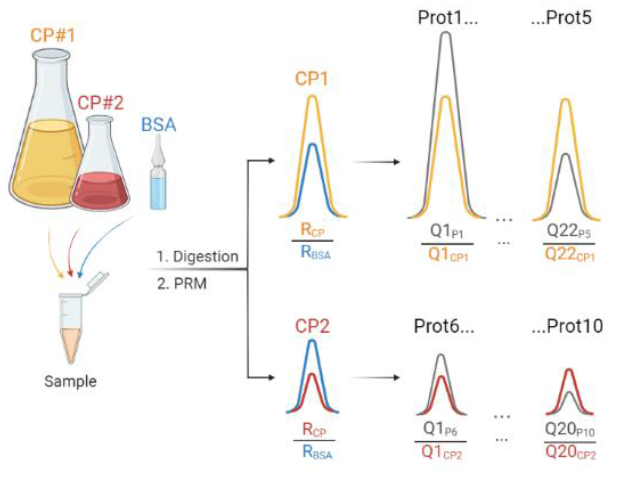

