## Supplementary material for "FastCAT accelerates absolute quantification of proteins by using multiple short non-purified chimeric standards": FastCAT_Supporting Information

Table of Content

| | |
|---|---|
| Figure S1 | Scheme of chimeric protein (CP) |
| Figure S2 | SDS-PAGE gel images of short CPs |
| Figure S3 | *E. coli* background in non-purified CP |
| Figure S4 | Amino acid sequences of CP05/CP06 |
| Figure S5 | Truncation patterns for CP05/CP06 |
| Figure S6 | Tandem mass spectra for $R_n$/$R_s$-peptides |
| Figure S7 | Relative $R_n$/$R_s$-peptides abundances for CP05/CP06 vs. BSA |
| Figure S8 | Precision of FastCAT method |
| Figure S9 | PCA plot for CSF proteins from patients with multiple sclerosis |
| Table S1 | *E. coli* background proteins used for comparison in Figure S2 |
| Table S2 | Error in quantification of CPs by $R_s$-peptides |
| Table S3 | Protein concentrations in CSF from individual patients |
| Table S4 | Amino acid sequences for all used CPs |

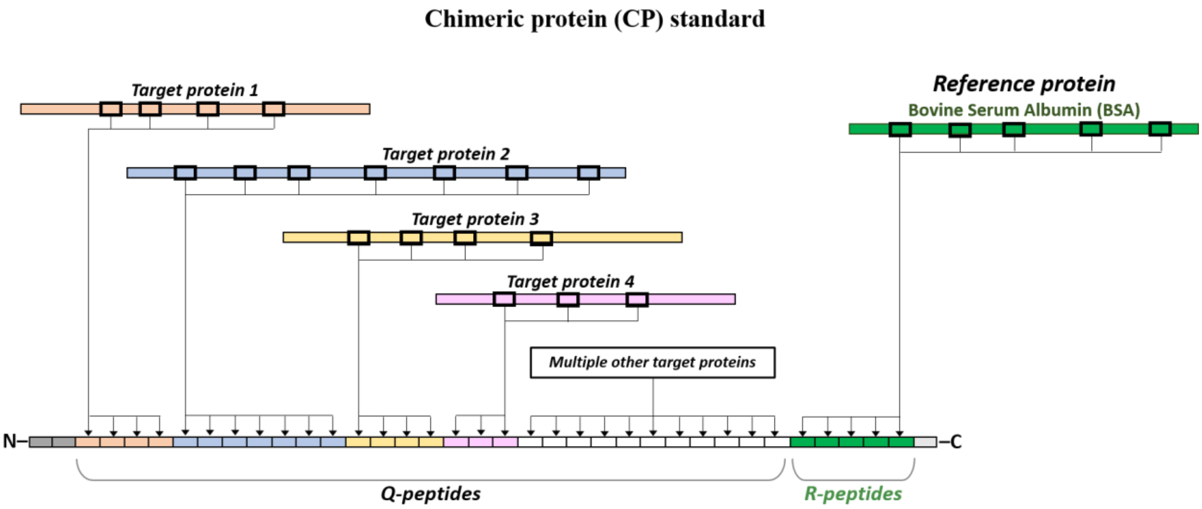


**Figure S1.** Scheme of chimeric protein (CP) standards used in this work. A typical CP comprises quantotypic Q-peptides selected from the sequences of target proteins and reference R-peptides selected from bovine serum albumin (BSA). They were concatenated *in-silico* into a chimeric sequence, whose N- and C-termini contain additional tag sequences (in gray) for R- and Q- peptides protection and (optionally) for the affinity purification.

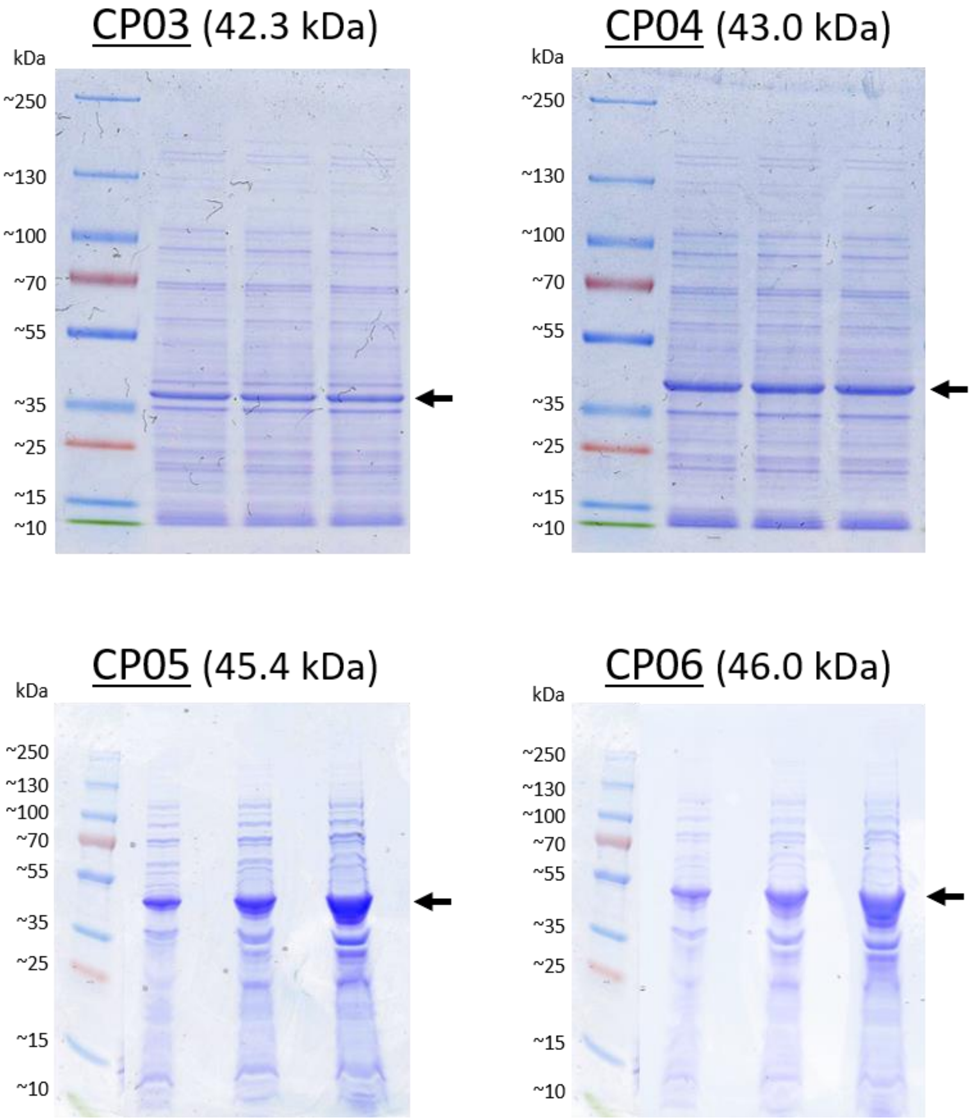


**Figure S2.** Images of Coomassie stained SDS-PAGE gels of four short (<50 kDa) CPs used in this study. Gel bands corresponding to the full-length CPs are designated with arrows.

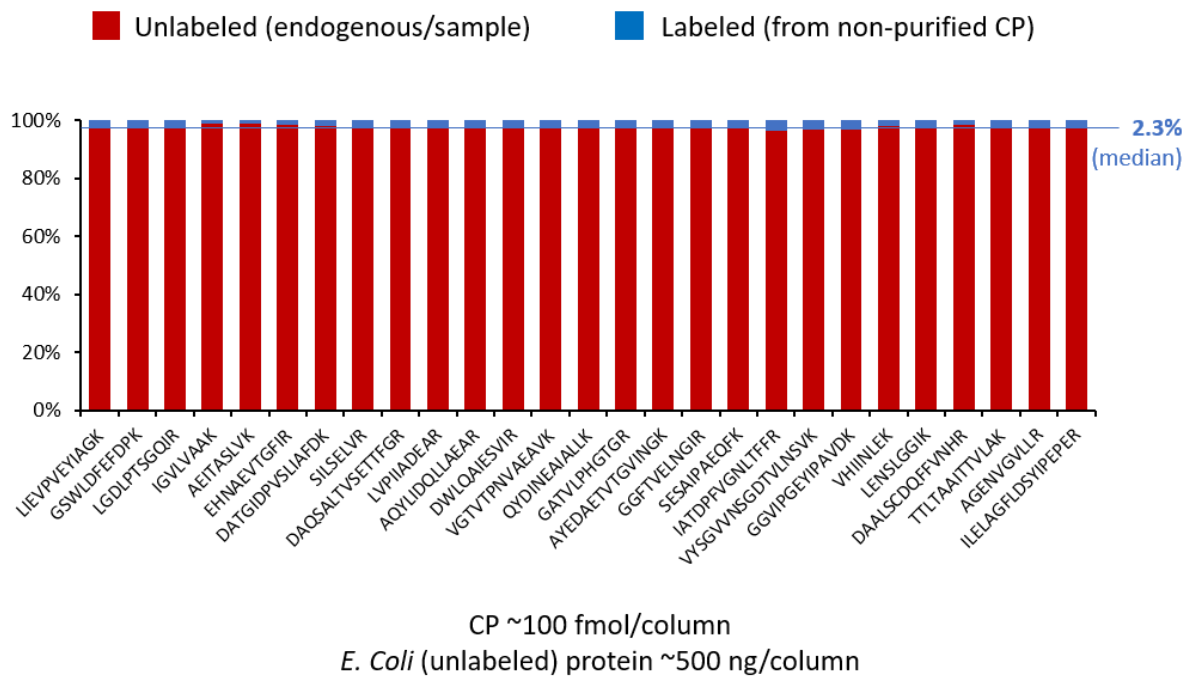


**Figure S3.** Contribution of extra background from the expression host (*E. coli*) introduced by spiking a unpurified CP standard (labeled, blue bars) compared to unlabeled *E. coli* proteins (unlabeled/endogenous, red bars). The protein amounts are typical for FastCAT experiments with CP at approximately 100 fmol and total protein of 500 ng loaded on column. The contribution of background (labeled) is ~2 % of the total protein. Nine abundant *E. coli* proteins (mainly ribosomal proteins and elongation factors) were considered. Their quantification was based on the abundance of at least three peptides having consistent L/H ratios (see Table S1). These peptides were detected as predominantly doubly charged ions, did not have missed cleavage sites and cysteine and methionine residues. Moreover, CP labeling efficiency exceeded 99 % and therefore all unlabeled peptides originated from background *E. coli* proteins.

**A** CP05

MGSAWSHPQFEKGGGSGGGSGGSAWSHPQFEKLEVLFQGPAAAKVFADYEEYVKDFYELEPHKV
AAAFPGDVDRGLAGVENVTELKHLVDEPQNLIKLVQETLEFAKSELEEQLTPVAEETRLGPLVEQGR
AATVGSLAGQPLQERQWAGLVEKLAVYQAGARNILTSNNIDVKIPTTFENGRNPNLPPETVDSLKV
LNQELRWYEIEKAEFVEVTKITLLSALVETRNLAVSQVVHKAVLDVFEEGTEASAATAVKADLSGITG
ARHVEDVPAFQALGSLNDLQFFRYSLTYIYTGLSKEIPAWVPFDPAAQITKQKWEAEPVYVQRNYPS
LELDKNYLNYGEEGAPGKGEAGAPGEEDIQGPTKWQQQGDLQDTKLGEYGFQNALIVRDAFIGTF
LYEYSRGSGHHHHHH

Apolipoprotein E (APOE) – Apolipoprotein D (APOD) – Alpha-1-antichymotrypsin (AACT) – Zinc-alpha-2-glycoprotein (ZAG) – Secretogranin-1 (SCG1)
BSA (R-)peptides: native and scrambled

**B** CP06

MGSAWSHPQFEKGGGSGGGSGGSAWSHPQFEKLEVLFQGPAAAKVFADYEEYVKDFYELEPHKV
AAAFPGDVDRGLAGVENVTELKHLVEEPNQLIKLVNELTEFAKVAAGAFQGLRGQTLLAVAKTLDLG
ENQLETLPPDLLRENQLEVLEVSWLHGLKFPLTNAIKILGQQVPYATKTLLSVGGWNFGSQRVTIDS
SYDIAKGNQWVGYDDQESVKADFVETVKFIPLIPIPERASPFPVYKTDGAAPNVAPSDVGGGGR
VQVTSQEYSARNSEPQDEGELFQGVDPRTHLGEALAPLSKLADLASDLLLQYLLQGGARVGEEDEE
AAEAEAEAEEAERFQLTFPLRTPAAETLSQLGQTLQSLKALSGNVIAWAESHIEIYGGATKLGDYGFN
NALIVRDAFLGSFLYEYSRGSGHHHHHH

Leucine-rich alpha-2-glycoprotein (A2GL) – Chitinase-3-like protein (CH3L1) – Contactin-1 (CNTN1) – Neurosecretory protein VGF (VGF) – Neuronal pentraxin-1 (NPTX1)
BSA (R-)peptides: native and scrambled

**Figure S4.** Amino acid sequences of the CP05 and CP06 constructs used for the quantification of 10 neurological protein markers in CSF samples.

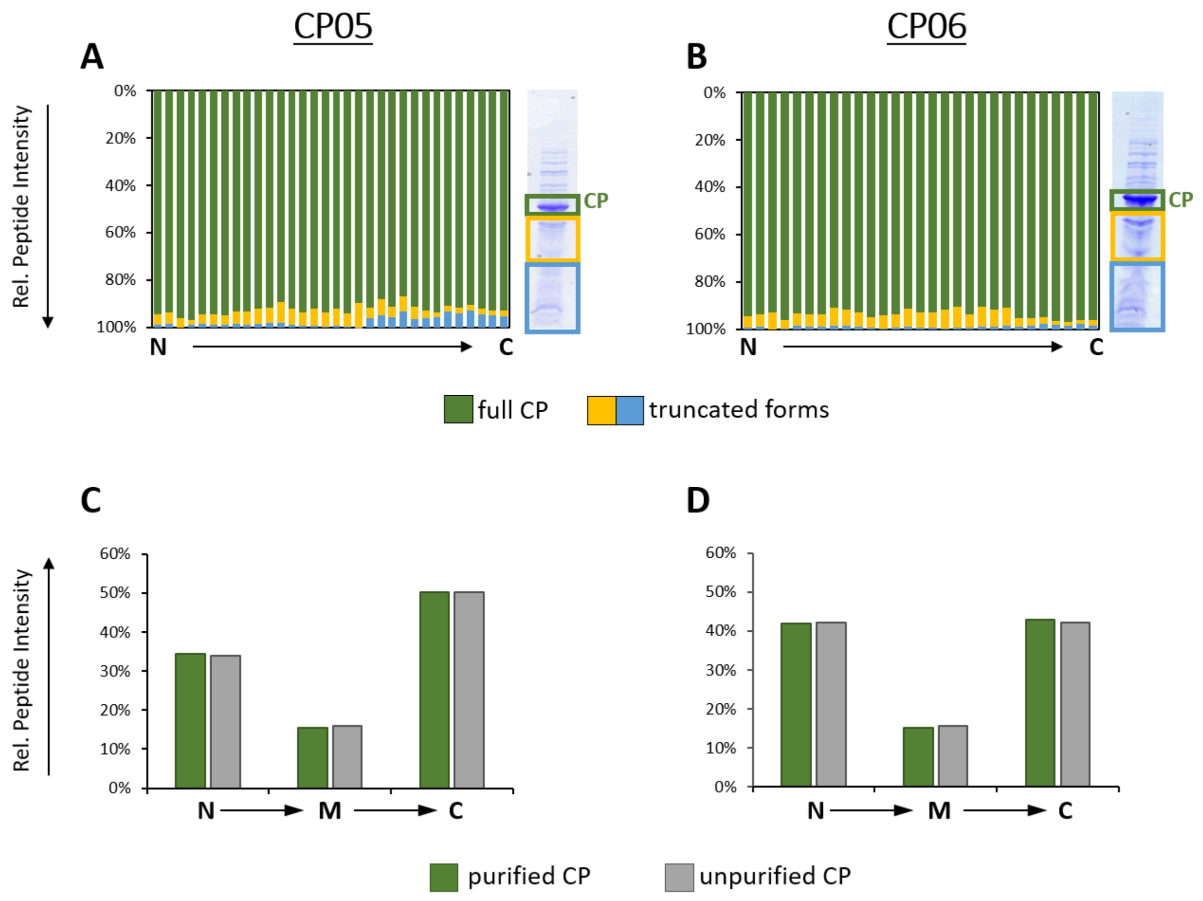


**Figure S5.** Truncation patterns for CP05 (A and C) and CP06 (B and D) standards.

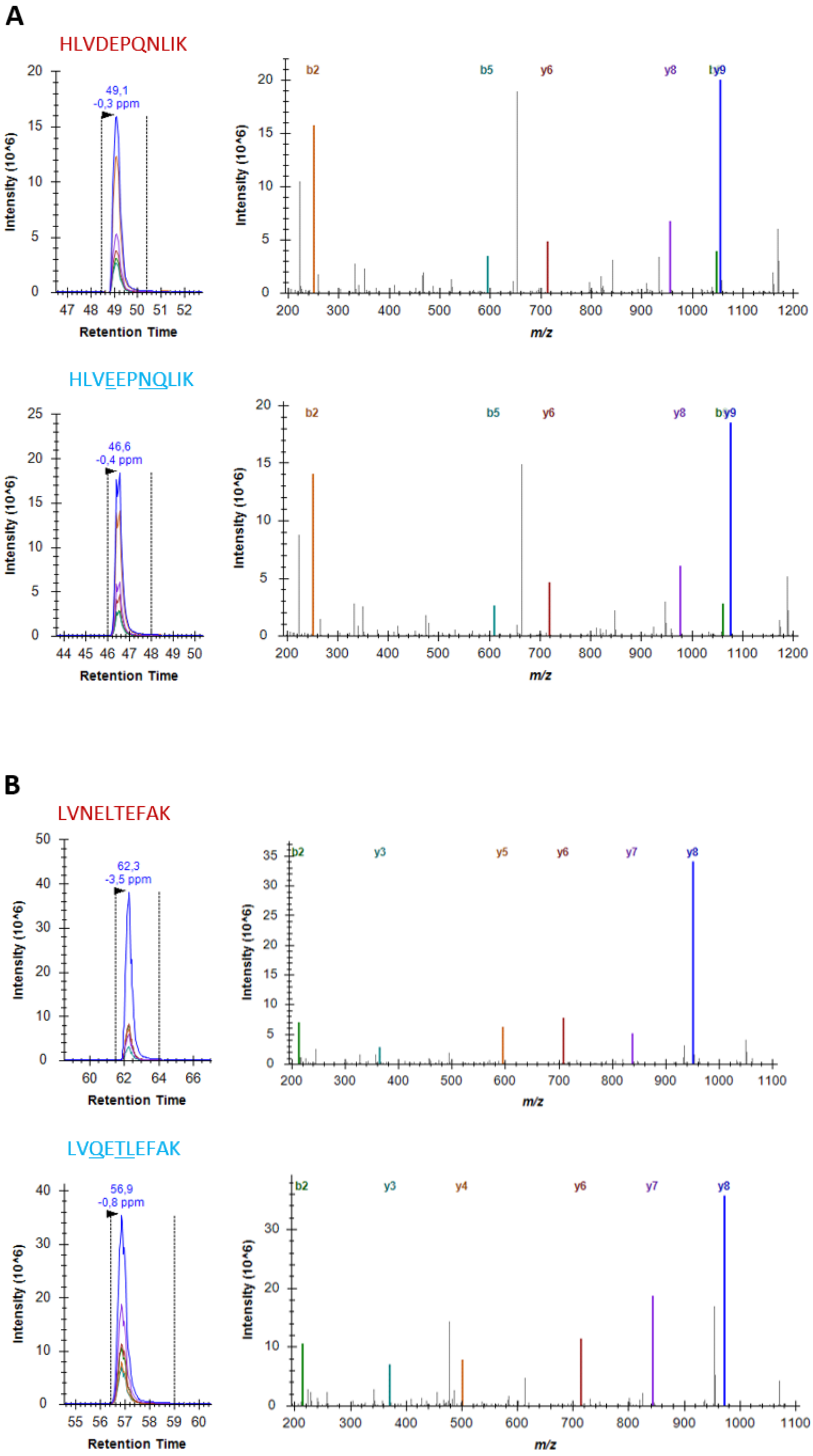
A
HLVDEPQNLIK
49,1
-0,3 ppm
Intensity (10^6)
Retention Time
b2
b5
y6
y8
y9
m/z
HLVEEPNQLIK
46,6
-0,4 ppm
Intensity (10^6)
Retention Time
b2
b5
y6
y8
y9
m/z
B
LVNELTEFAK
62,3
-3,5 ppm
Intensity (10^6)
Retention Time
b2
y3
y5
y6
y7
y8
m/z
LVQETLEFAK
56,9
-0,8 ppm
Intensity (10^6)
Retention Time
b2
y3
y4
y6
y7
y8
m/z

C

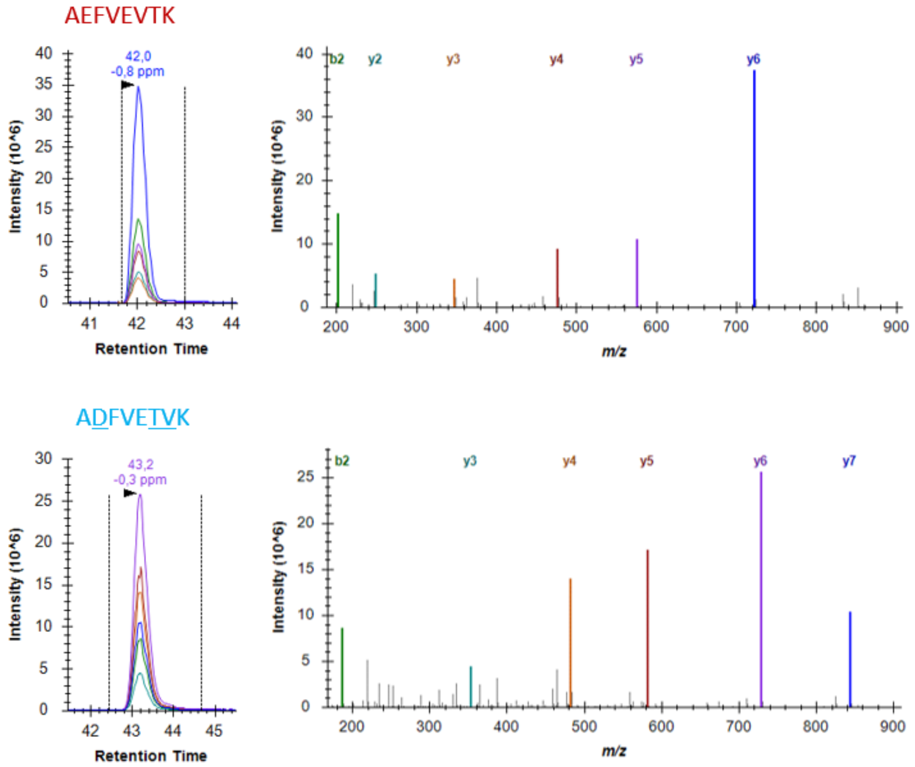
AEFVEVTK
42,0
-0,8 ppm
Intensity (10^6)
Retention Time
b2
y2
y3
y4
y5
y6
m/z
ADFVETVK
43,2
-0,3 ppm
b2
y3
y4
y5
y6
y7

D

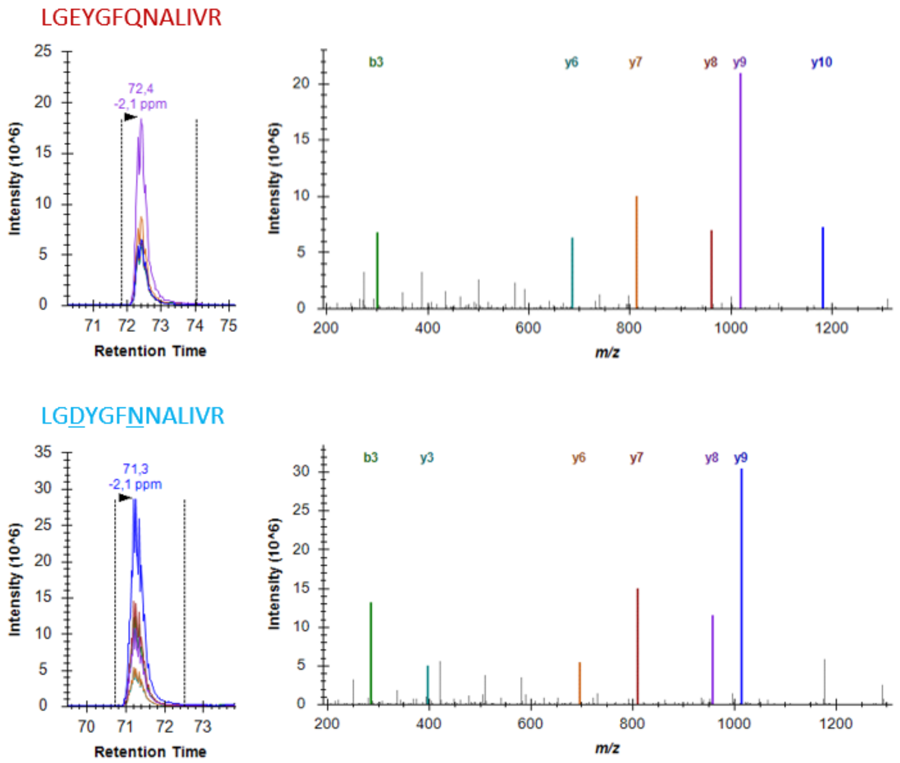
LGEYGFQNALIVR
72,4
-2,1 ppm
Intensity (10^6)
Retention Time
b3
y6
y7
y8
y9
y10
m/z
LGDYGFNNALIVR
71,3
-2,1 ppm
b3
y3
y6
y7
y8
y9

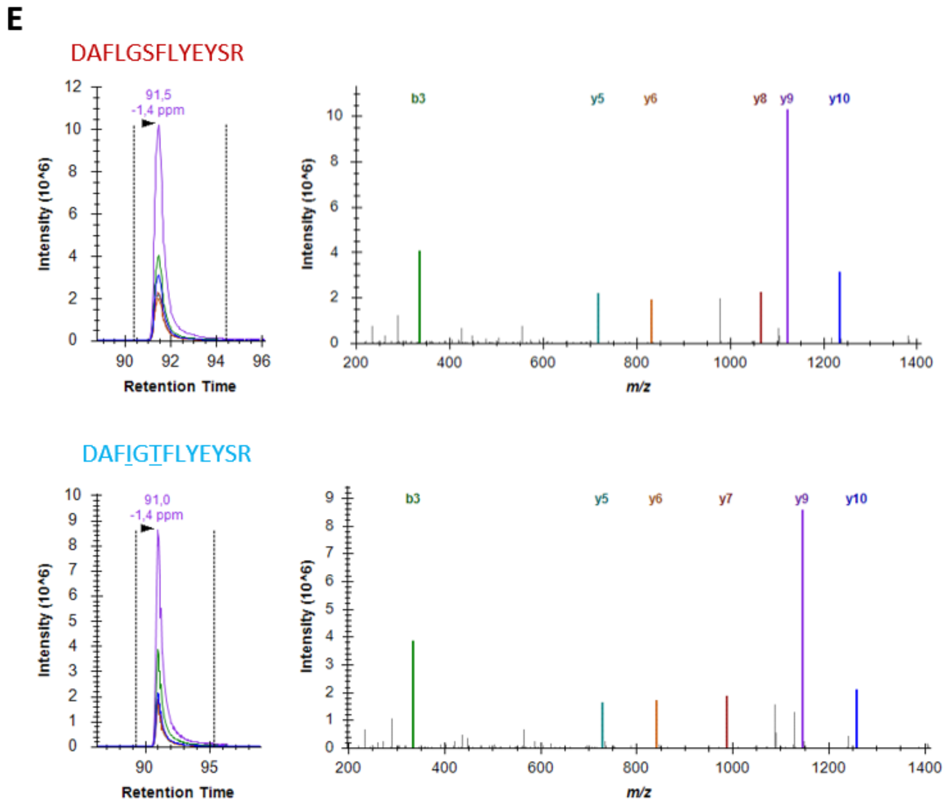


**Figure S6.** Tandem mass spectra of R-peptides in native (BSA, unlabeled) and scrambled forms (CP05/06, labeled) together with their chromatographic peaks: HLVDEPQNLIK (nat) and HLVEEPNQLIK (scr) (**A**), LVNELTEFAK (nat) and LVQETLEFAK (scr) (**B**), AEFVEVTK (nat) and ADFVETVK (scr) (**C**), LGEYGFQNALIVR (nat) and LGDYGFNNALIVR (scr) (**D**), DAFLGSFLYEYSR (nat) and DAFIGTFLYEYSR (scr) (**E**).

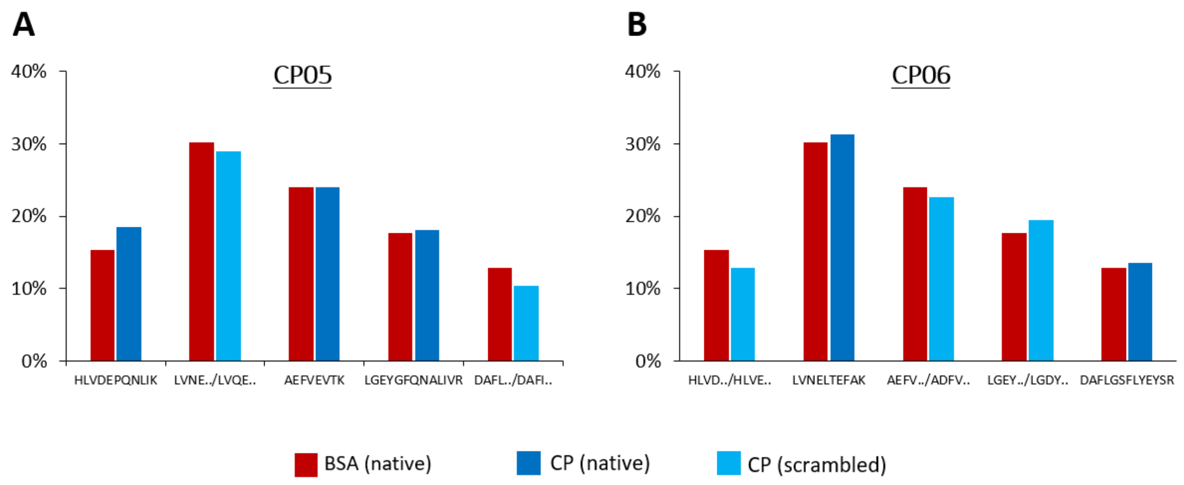


**Figure S7.** Matching relative peak areas of R-peptides for BSA (native) and both CP05 (**A**) and CP06 (**B**) obtained using Top6(y,b) fragment ions. Differences were <5 % for all pairs.

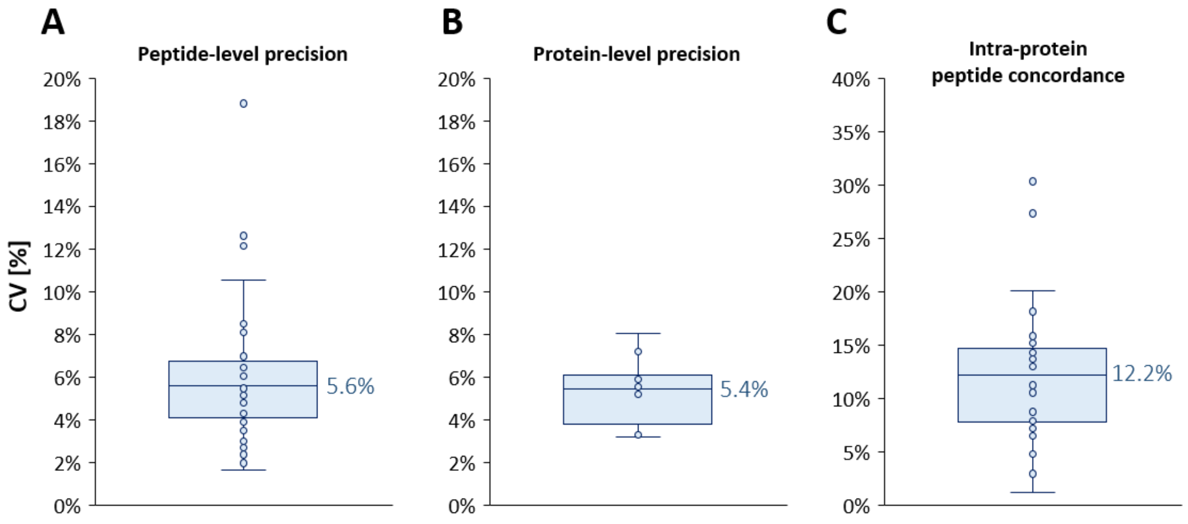


**Figure S8.** Precision of the FastCAT quantification estimated from the PRM analysis of the pooled CSF sample. Panels A and B show peptide-level and protein-level precision, respectively. The graphs are created with data acquired for all investigated proteins (10 target proteins and two CPs) based on three sample preparation replicates and two LC injections each. Panel C shows intra-protein peptide concordance obtained from the same data set. Median CV values are provided at the plots. Each box plot displays the median (line), the 25$^{th}$ and 75$^{th}$ percentiles (box), and the 5$^{th}$ and 95$^{th}$ percentiles (whiskers).

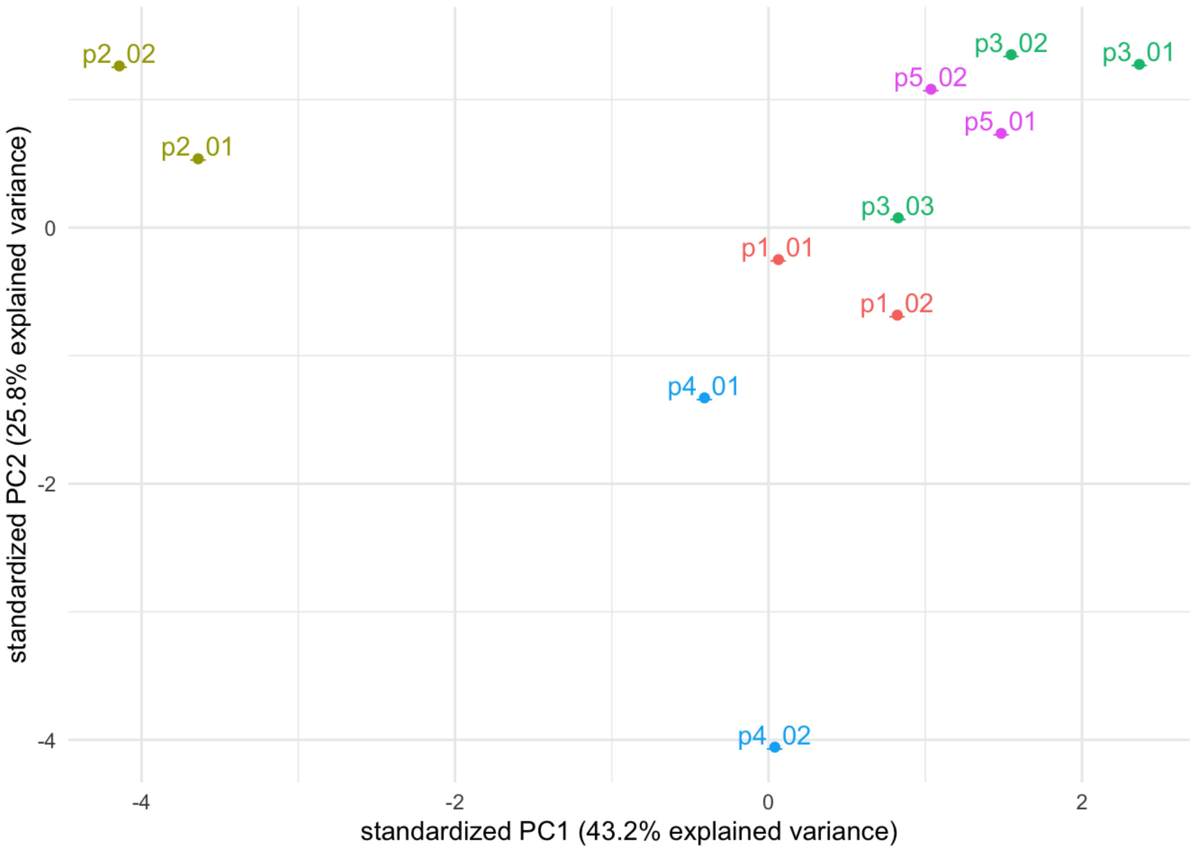


**Figure S9.** Principal component analysis (PCA) plot for all protein quantities (from Table S4) in patients 1-5 (p1-p5) with two (or three for p3) collections of samples. In this way, p2_01 and p2_02 stand for the first and second CSF collection from the patient p2, respectively.

**Table S1.** List of representative *E. coli* proteins (selected from most abundant proteins) used for estimating the amount of extra protein background introduced by spiking an unpurified CP standard, For each protein we provide UniProt accession number; sequences of peptides used for the quantification (three peptides per each protein); peptides retention times and charge states.

| Protein | UniProt Accession | Peptide | RT | Charge state |
|---|---|---|---|---|
| DNA-directed RNA polymerase | P0A8V2 | LIEVPVEYIAGK | 76.7 | 2 |
| | | GSWLDFEFDPK | 87.1 | 2 |
| | | LGDLPTSGQIR | 57.5 | 2 |
| Elongation factor Ts | P0A6P1 | IGVLVAAK | 53.2 | 2 |
| | | AEITASLVK | 55.2 | 2 |
| | | EHNAEVTGFIR | 53.7 | 3 |
| 50S ribosomal protein L4 | P60723 | DATGIDPVSLIAFDK | 88.9 | 2 |
| | | SILSELVR | 76.7 | 2 |
| | | DAQSALTVSETTFGR | 69.9 | 2 |
| Pyruvate dehydrogenase E1 | P0AFG8 | LVPIIADEAR | 64.4 | 2 |
| | | AQYLIDQLLAEAR | 89.5 | 2 |
| | | DWLQAIESVIR | 105.7 | 2 |
| 50S ribosomal protein L1 | P0A7L0 | VGTVTPNVAEAVK | 56.0 | 2 |
| | | QYDINEAIALLK | 85.9 | 2 |
| | | GATVLPHGTGR | 41.1 | 2 |
| 30S ribosomal protein S1 | P0AG67 | AYEDAETVTGVINGK | 64.3 | 2 |
| | | GGFTVELNGIR | 70.9 | 2 |
| | | SESAIPAEQFK | 55.2 | 2 |
| Elongation factor G | P0A6M8 | IATDPFVGNLTFFR | 89.9 | 2 |
| | | VYSGVVNSGDTVLNSVK | 67.9 | 2 |
| | | GGVIPGEYIPAVDK | 70.7 | 2 |
| 30S ribosomal protein S2 | P0A7V0 | VHIINLEK | 51.9 | 2 |
| | | LENSLGGIK | 51.7 | 2 |
| | | DAALSCDQFFVNHR | 66.1 | 3 |
| Elongation factor Tu1 | P0CE47 | TTLTAAITTVLAK | 82.1 | 2 |
| | | AGENVGVLLR | 61.4 | 2 |
| | | ILELAGFLDSYIPEPER | 98.4 | 2 |

**Table S2.** Relative error of the CP05/06 quantification based on the scrambled R-peptides and using either MS1-derived (DDA) peptides peak areas or different MS2 fragment ions (PRM).

| Native form | Scrambled form in CP | Δ RT [min] | CP with scrambled form | MS1-based (DDA) | | | MS2-based (PRM) | | | | | | |
|---|---|---|---|---|---|---|---|---|---|---|---|---|---|
| | | | | CP [fmol] by nat | CP [fmol] by scr | % error | CP [fmol] by nat | CP [fmol] by scr | | | | | |
| | | | | | | | | Top1 | % error | Top3(y) | % error | Top6(y,b) | % error |
| HLVDEPQNLIK | HLVEEPNQLIK | 2.5 | CP06 | 275 | 290 | 5.4% | 294 | 241 | 17.9% | 241 | 17.7% | 236 | 19.7% |
| LVNELTEFAK | LVQETLEFAK | 5.4 | CP05 | 270 | 295 | 9.0% | 280 | 180 | 35.8% | 245 | 12.5% | 250 | 10.9% |
| AEFVEVTK | ADFVETVK | 1.2 | CP06 | 275 | 274 | 0.4% | 294 | 185 | 36.9% | 268 | 8.7% | 265 | 9.7% |
| LGEYGFQNALIVR | LGDYGFNNALIVR | 1.1 | CP06 | 275 | 312 | 13.4% | 294 | 329 | 12.1% | 341 | 16.4% | 310 | 5.4% |
| DAFLGSFLYEYSR | DAFIGTFLYEYSR | 0.5 | CP05 | 270 | 188 | 31.0% | 280 | 214 | 23.9% | 203 | 27.4% | 211 | 24.7% |

**Table S3**. Protein concentrations (ng/mL) in 11 CSF samples from 5 patients diagnosed with multiple sclerosis. CSF was collected twice for four patients and three times for the patient #3. Patient age during the 1st collection is given in years (y), while the time difference (Δt) to the 2nd (and 3rd for patient 3) puncture is given in months (m).

| **Protein** | **Concentration in CSF [ng/mL]** | | | | | | | | | | |
|---|---|---|---|---|---|---|---|---|---|---|---|
| | **Patient 1** | | **Patient 2** | | **Patient 3** | | | **Patient 4** | | **Patient 5** | |
| | time 1 | time 2 | time 1 | time 2 | time 1 | time 2 | time 3 | time 1 | time 2 | time 1 | time 2 |
| | age, 43 y | Δt, 40 m | age, 61y | Δt, 17 m | age, 36 | Δt, 10 m | Δt, 16 m | age, 30 y | Δt, 32 m | age, 30 y | Δt, 26 m |
| **APOE** | 1489.28 | 1604.15 | 1106.19 | 922.07 | 1926.15 | 1807.81 | 1772.67 | 1854.96 | 2075.59 | 1996.34 | 1811.10 |
| **APOD** | 1197.68 | 1307.78 | 1185.15 | 1203.23 | 1284.59 | 1281.67 | 1297.31 | 1083.58 | 1060.41 | 1246.55 | 1327.29 |
| **AACT** | 759.84 | 684.61 | 765.58 | 723.96 | 580.30 | 564.96 | 770.17 | 810.35 | 1127.28 | 540.90 | 588.94 |
| **ZAG** | 100.05 | 113.18 | 102.01 | 96.32 | 85.98 | 85.89 | 87.58 | 102.12 | 108.05 | 85.19 | 85.76 |
| **SCG1** | 636.56 | 684.09 | 451.77 | 388.55 | 942.32 | 885.69 | 869.22 | 788.86 | 862.85 | 852.69 | 850.08 |
| **LRG1** | 67.98 | 71.42 | 77.14 | 80.84 | 50.02 | 47.78 | 114.29 | 38.45 | 92.91 | 27.13 | 33.82 |
| **CH3L1** | 51.77 | 35.09 | 35.16 | 33.37 | 53.82 | 49.85 | 44.63 | 38.90 | 27.48 | 27.90 | 25.74 |
| **CNTN1** | 238.90 | 256.07 | 159.70 | 148.28 | 229.68 | 230.81 | 229.77 | 241.20 | 268.62 | 257.04 | 246.15 |
| **VGF** | 139.34 | 163.29 | 62.64 | 60.65 | 175.94 | 133.49 | 141.99 | 73.35 | 98.14 | 110.97 | 99.89 |
| **NPTX1** | 75.67 | 86.02 | 42.50 | 37.18 | 77.01 | 72.52 | 71.98 | 64.28 | 81.85 | 70.87 | 68.53 |

**Table S4.** Chimeric protein (CP) standards used in this study together with their molecular weights, labeling efficiencies as well as amino acid sequences.

| Chimeric Protein | Molecular weight | Labeling efficiency | Amino acid sequence |
|---|---|---|---|
| CP01 | 264.8 kDa | >99 % | MGSAWSHPQFEKGGGSGGGSGGSAWSHPQFEKLEVLFQGPAAAKVFADYEEYVKDFYELEPHKVAAAFPGD<br>VDRGLAGVENVTELKGSTLSAQQGSQFKNEPVSYILELINGRNIGDSYVAAKLDNGYNHLIEVVRLVLGIPTYG<br>RFSPLVASNERLTEAEGSSLYIGGRSTDAEEDPQVIKFADQDNDLVNLRLSNFVDTTVAWLNYRFFVSVTRVQL<br>TDLNRDVFIPDVFNNYKILAIFPFPGPSQYINVVPYLKVPIWPYTLYFRGTFFVSHKGHEISVFSLTHKAPLGLHF<br>HASWLKIINNPEATQRAVYWVEHVSRSHYHVGSALAKAEQNGYGVTVHYEELSSAKGSAYAHAENTLRSEGD<br>LTQITPPSALGVQVRAEVEAVQIIAESLKAANQLTDRPTIINVAISPSSDRGHGAGGADVLTYKDLNQGNSYVQD<br>KSLVFVDNHDNQRSEYTGLGAITEFRVVEFLDHLIDLGVAGFRIGALDTSRLIELLVRLADLLLQEERVNGEPVD<br>VGAFFIRITPDVVGAIVQEATTYFRATQLSEQLGSELTPKLLAADYADGVSQPRILSWNAVNLYGLKASEQPGL<br>TAIHTAFLRLLPAQYEDGISAPRGIATNDVGIAKDTFSFLTGSGRAINEGGFQSLPQKFGLNEGSEPQAYGIGLKI<br>TTHYTLNPRAILVYLVEKITPGWEENWAGALDVKINPQHSIPTLVDNGFTIWESRALGLEFNKINLDPYALYVIV<br>EKTNFDDSALFYASGESLKVLTTGGDAGSNRLGSDVQPPGRVFLQLLENQRNLSQTFGNIWRVFDLPDFLTWR<br>GSTVQFAARDGAISHVVFKYFNAYLSDRFSEPYDSTLSDVIKNVANLEATKDADAIVQQTLAKGTQAEIVVAD<br>VTKVGDVTEVAEAVAFLASSKSVLLTLRIDALEGVYKALNLDFDYKAITFPLFWENKLTLYGIDGSPPVRELPY<br>YEEANGSRAVGVELNKAPADPEAFKAIQVYLVEKLLNLQAGEHLKPEFLKINPQHTIPTLVDNGFALWESRGL<br>VVSGTRSLSDVSLTGRGNSYLILPPPRISGPSNHVTVVRNAAFSGDSYVSHRSVVNVGLGQALRSTQEEVDHIRS<br>AVHDVEVFLKVSTAIDAISVSGRVALQIQDVATSSRLEEISLRLLELHDNRLFNNFDVLRLAVLDLSHNRASLSG<br>IQSHAFKTFFDGNPIHTLRDFGVELEDLQITRSTISSTTTVRTSTTSSLTGNPRTSVATVAGGAVVGATKLLVPGS<br>SSTTTTSSLRLLPDTTDEQLLTSALEEKSTGNIFAAKVFAVNSAGRIFADNVYGRGVAEDFAPSFVKHDGGSPIT<br>GYIIEKHYPNPAVRVVGSEADTGRDAPTTESYLRSTVPFATSESNRLVGGNDLDTPLIITRIAELEGLGSRIQYLE<br>DSTRIGEAISQSSIRFITQIVDVTKAEDAGVYLVVARTYYDFGFVALDIIKVWQVSVSAHALLWDLNDGKDVLS<br>VAFSADNRLWDLAAGKLNNDLIARFQEALAGLSKLAAHDALGGAAKSAGSGVSTTAIEKIVQVQIDDVGKSEL<br>DVFSDWLQVARFSQSDFGLDQGETLLRESLLEITIYHQKGAAYQEAPVDAEVAVTPKISAFGLFTYSVFSIIAGS<br>LKVFANPVQLEFYGFVPWSIIQRQVGASSGSSGAVSKIVQIQQVGPAHLNVKVQDNSVLVEGNHEERVLPAIEA<br>FQKFSEATLDEIIRFHGVALAFNALDSKENAILTDIWNITPFKVIVDILLKTILVDLQVGKLAELHAASVVAKEAG<br>LEIELAPKAGFAGDDAPRSYELPDGQVITIGNERVAPEEHPVLLTEAPLNPKGYSFTTTAEREITSLAPSTIKEITA<br>LAPSTIKQEYDESGPGIVHRQEYDESGPSIVHRSIELLEKLQASGDFPTTKEQNSPIYISRATVAAFAASEGHAHPR<br>YQVIVKSSDAQSQATASEAESKALSASSLIALSSRDPVLTAFQLSWELKDDYGITEDDIIEVRATAWVIVHRVEL<br>ELWLKVVLPDLAVLRALGIEEQSGGAARVDEYGFFLYWKYYLSDLARIEQADYLPTEQDILRVPTTGILEYPFD<br>LDGIVFRTIITYPWFQNSSVILFLNKAGIAVEGDIKDAFGIIVSYAVKDTALASTTLIASQDARDFLLSPGELELEV<br>TLDKVFGQLATTYRLQYAPLNRISLEVTLDRLLQPAPGTIEKFYPEDVAEELIQDITLRIGFPWSEIRTTHTAGFL<br>ANDRLFYLQVKAAIELNRGDNILLTKVPPEVPPSKLVDFIINRSLVGVQKQIDVETVADRYIQATDLSEASRNAE<br>YDQFLEDIQKATAWVIVHRALSASSLIALSSRHLVDEPQNLIKYLYEIARQTALVELLKLGEYGFQNALIVRDAF<br>LGSFLYEYSRLVNELTEFAKGSGHHHHHH |

| | | | |
|---|---|---|---|
| CP02 | 79.3 kDa | >99 % | MGSAWSHPQFEKGGGSGGGSGGSAWSHPQFEKLEVLFQGPAAAKVFADYEEYVKDFYELEPHKVAAAFPGD<br>VDRGLAGVENVTELKLGDYGFNNALIVRDAFIGTFLYEYSRAAAYVLQETPVVNALVDENEIVYRSDGEVEIA<br>SEKVIDTAYEIIKGIILIGEGIGNAEEQAAEFLKVQNDDSIVFFDYRVLEATLAQDFSKLVTWYDNEFGYTNRYA<br>GEDAAAGAAETLFVAKAEIEAVQIIAETLKVVEFLEHILDLGVAGFRIVQVNLDDVGKVGDVTEVAEAAVFLA<br>TSKAGLAIEGDIKANEILSDIWNITPFKGIAEDFAPTFVKIADLEGIYKALQLDFEYKALVFETWQGPLEVRWVAI<br>DGEQYGEGSSRLIDDNVANALKVVDHAYEAVVIGAGGAGLRYLYDVARFHGATSINLVGDLDTVTNPKGTVA<br>HDGDYLIVAKTVEADAAHGSVTRTIEADAAHGSVTRVVELITYIATKYAVFDTGSRVTENVLAFIYKQLLFSAG<br>AELNKLDLGTVVSPVSGPKLGANTLLELVIFGRAFGGNTQDFGRVFQFLEASAGSKLNADTSLFILASKGQETST<br>QPIATIFAWSRAGQSHLGLPIFGSAVEAKGVEPSHAISGARALIANGTGPYFYLPKNTVIASGGYGRAAAAQINY<br>IRSGNVVPGYHGAVLRVALLGAGAGIGNPLGLLLKVPQVILAVGLPARHLVEEPNQLIKLVNELTEFAKGSGHH<br>HHHH |
| CP03 | 42.3 kDa | >99 % | MGSAWSHPQFEKGGGSGGGSGGSAWSHPQFEKLEVLFQGPAAAKVFADYEEYVKDFYELEPHKVAAAFPGD<br>VDRGLAGVENVTELKIGEEYISDLDQLRVIFLENYRLLSYVDDEAFIRVLVDLERTLSDYNIQKESTLHLVLRTIT<br>LEVEPSDTIENVKEGIPPDQQRIGDYAGIKVLGIDGGEGKEALDFFARVVGLSTLPEIYEKANELLINVKGVIFYE<br>SHGKGNPTVEVELTTEKTFAEALRAADALLLKTAGIQIVADDLTVTNPKNVNDVIAPAFVKVNQIGTLSESIKA<br>VDDFLISLDGTANKHLVDEPQNLIKYLYEIARQTALVELLKLGEYGFQNALIVRDAFLGSFLYEYSRLVNELTEF<br>AKGSGHHHHHH |
| CP04 | 43.0 kDa | >99 % | MGSAWSHPQFEKGGGSGGGSGGSAWSHPQFEKLEVLFQGPAAAKVFADYEEYVKDFYELEPHKVAAAFPGD<br>VDRGLAGVENVTELKHLVDEPQNLIKLVNELTEFAKIGEEYISDLDQLRVIFLENYRLLSYVDDEAFIRVLVDLE<br>RTLSDYNIQKESTLHLVLRTITLEVEPSDTIENVKEGIPPDQQRYLYEIARLVTDLTKAEFVEVTKIGDYAGIKVL<br>GIDGGEGKEALDFFARVVGLSTLPEIYEKANELLINVKGVIFYESHGKGNPTVEVELTTEKTFAEALRAADALLL<br>KTAGIQIVADDLTVTNPKNVNDVIAPAFVKVNQIGTLSESIKAVDDFLISLDGTANKLGEYGFQNALIVRDAFLG<br>SFLYEYSRGSGHHHHHH |
| CP05 | 45.4 kDa | >99 % | MGSAWSHPQFEKGGGSGGGSGGSAWSHPQFEKLEVLFQGPAAAKVFADYEEYVKDFYELEPHKVAAAFPGD<br>VDRGLAGVENVTELKHLVDEPQNLIKLVQETLEFAKSELEEQLTPVAEETRLGPLVEQGRAATVGSLAGQPLQ<br>ERQWAGLVEKLAVYQAGARNILTSNNIDVKIPTTFENGRNPNLPPETVDSLKVLNQELRWYEIEKAEFVEVTKI<br>TLLSALVETRNLAVSQVVHKAVLDVFEEGTEASAATAVKADLSGITGARHVEDVPAFQALGSLNDLQFFRYSL<br>TYIYTGLSKEIPAWVPFDPAAQITKQKWEAEPVYVQRNYPSLELDKNYLNYGEEGAPGKGEAGAPGEEDIQGP<br>TKWQQQGDLQDTKLGEYGFQNALIVRDAFIGTFLYEYSRGSGHHHHHH |
| CP06 | 46.0 kDa | >99 % | MGSAWSHPQFEKGGGSGGGSGGSAWSHPQFEKLEVLFQGPAAAKVFADYEEYVKDFYELEPHKVAAAFPGD<br>VDRGLAGVENVTELKHLVEEPNQLIKLVNELTEFAKVAAGAFQGLRGQTLLAVAKTLDLGENQLETLPPDLLR<br>ENQLEVLEVSWLHGLKFPLTNAIKILGQQVPYATKTLLSVGGWNFGSQRVTIDSSYDIAKGNQWVGYDDQESV<br>KADFVETVKFIPLIPIPERASPFPVYKTDGAAPNVAPSDVGGGGGRVQVTSQEYSARNSEPQDEGELFQGVDPRT<br>HLGEALAPLSKLADLASDLLLQYLLQGGARVGEEDEEAAEAEAEAEEAERFQLTFPLRTPAAETLSQLGQTLQS<br>LKALSGNVIAWAESHIEIYGGATKLGDYGFNNALIVRDAFLGSFLYEYSRGSGHHHHHH |
